# SPEECHLESS duplication in grasses expands potential for environmental regulation of stomatal development

**DOI:** 10.1101/2025.07.29.667563

**Authors:** Joel M. Erberich, Britney Bennett, Dominique C. Bergmann

## Abstract

Plants regulate stomatal development and function to acquire atmospheric carbon dioxide for photosynthesis while minimizing water loss. The ancestral basic helix-loop-helix transcription factor (TF) gene that drove stomata production in early land plants diversified to become paralogs *SPEECHLESS* (*SPCH*), *MUTE*, and *FAMA*. Extant grasses exhibit a particularly interesting set of duplications and losses of *SPCH*.

Using phylogenetic methods, we tracked the history of SPCH duplications. *Brachypodium distachyon* and *Oryza sativa* plants bearing mutations in either *SPCH1* or *SPCH2, and B. distachyon* plants with SPCH1 or SPCH2 translational reporters were assayed under different environmental conditions for their effects on stomatal development.

We identified the Poaceae-specific *rho* whole genome duplication as the origin of *SPCH1* and *SPCH2* and demonstrated that both paralogs remain under selection. We found paralog-specific divergence in response to two environmental perturbations in both *B. distachyon and O. sativa*. Plausible molecular mechanisms underpinning paralog divergence, and cellular mechanisms driving the stomatal phenotypes are supported by analyses of *BdSPCH1* and *BdSPCH2* RNA and protein expression and by sequence variation among grasses.

These studies suggest ways in which a duplication of a key stomatal regulator enables adaptation and could inform genetic strategies to mitigate anticipated stressors in agronomically important plants.

## Introduction

Stomata are cellular complexes that enable efficient gas exchange between the aerial portions of plants and the environment. Stomata appeared in the fossil record coincident with the appearance of land plants and, along with the waxy cuticle, are thought to enable plants’ success on land (Payne, 1979). Among most clades of living and fossil plants, stomatal morphology is simple–two epidermal guard cells flanking a central pore. The morphology of neighboring cells and patterns of stomatal distribution on leaves vary considerably, however, and are useful taxonomic characters (Bergmann and Sack, 2007). Based on anatomical observations of preserved samples, it has been hypothesized that the developmental pathways to create stomata have increased in complexity over evolutionary time (Rudall *et al*., 2013). In parallel, genes encoding the core fate-regulating transcription factors have also expanded. Phylogenetic studies link the origin, duplication and diversification of subfamily Ia and subfamily IIIb basic helix-loop-helix (bHLH) transcription factors (TFs) to stomatal lineage diversity (Chen *et al*., 2017; Clark *et al*., 2022; Harris *et al*., 2020; Pires and Dolan, 2010). Genetic studies in the bryophyte moss *Physcomitrium patens* found a single subfamily Ia and subfamily IIIb pair are required to produce stomata at the base of the sporophyte spore capsule where they serve to promote dehiscence (Chater *et al*., 2016). In the dicot angiosperm *Arabidopsis thaliana*, stomata are the end product of a multi-step developmental pathway. Initiation, commitment, and differentiation steps in this pathway are each uniquely regulated by a different stage-specific subfamily Ia bHLH, SPEECHLESS (SPCH), MUTE, or FAMA, respectively (Bergmann *et al*., 2004; Ohashi-Ito and Bergmann, 2006; Pillitteri *et al*., 2008). Two Class IIIb factors, SCREAM (SCRM1) and SCRM2 are redundant heterodimer partners for all three subfamily Ia factors (Kanaoka *et al*., 2008). Orthologues of SPCH, MUTE, FAMA, SCRM1 and SCRM2 can be identified in many angiosperm genomes, and functional studies in *Solanum lycopersicum* (tomato), *Oryza sativa* (rice), *Brachypodium distachyon* (purple false brome) and *Zea mays* (maize) found these TFs to be generally conserved in their expression patterns and function in stomatal development (Liu *et al*., 2009; Nir *et al*., 2023; Ortega *et al*., 2019; Raissig *et al*., 2016; Wu *et al*., 2019). Intriguingly, novel stomatal forms redeploy the core bHLHs in new ways, for example grass stomata are dumbbell-shaped guard cells flanked by subsidiary cells, and MUTE has acquired new roles specifying the identity and position of both guard cells and subsidiary cells (Raissig *et al*., 2017; Spiegelhalder *et al*., 2024; Wu *et al*., 2019).

Stomatal function and development are subject in environmental regulation, and several studies have tracked the evolution of pathways enabling such regulation (Clark *et al*., 2022). When environmental response pathways have been described in molecular detail, their impacts on stomatal development typically involve SPCH. In *A. thaliana*, PHYTOCHROME INTERACTING FACTOR4 (PIF4) represses *SPCH* expression in warm temperatures (Lau *et al*., 2018) and mutations targeting the tomato *SPCH* promoter render plants unable to change stomatal production in response to light or temperature cues (Nir *et al*., 2023). Light and drought affect AtSPCH protein accumulation by modulating the signaling pathways whose downstream kinases phosphorylate AtSPCH (Kumari *et al*., 2014; Wang *et al*., 2021). That AtSPCH is the prime target of environmental regulation may not be surprising given its role as regulator of the asymmetric cell divisions that enable initiation and stem-cell like proliferation of stomatal lineages.

Grasses are an economically and ecologically important group of monocot angiosperms. They thrive in dry environments, in part due to their highly efficient stomata featuring dumbbell shaped guard cells and adjacent subsidiary cells (Franks and Farquhar, 2007; Leakey *et al*., 2019; Linder *et al*., 2018). Stomata are arranged in linear cell files in the leaf, with each stoma oriented in the same direction. The highly organized progression of stomatal formation begins with an asymmetric cell division, but unlike *A. thaliana*, there is no stem-cell like phase of repeated divisions, instead the smaller daughter of the asymmetric division is immediately destined to become a pair of guard cells. As in dicots, grass stomatal production can be modified by environmental conditions or precision genetic engineering, the latter of which has been shown to improve water-use-efficiency in a variety of agronomically important cereals (Caine *et al*., 2019; Dunn *et al*., 2019; Hughes *et al*., 2017; Mohammed *et al*., 2019). Whether SPCH is the target of environmental tuning in grasses is an open question and is particularly interesting because of (1) the more streamlined development in grass stomatal files and (2) because there are two SPCH paralogs in most grasses. The paralogs *BdSPCH1* and *BdSPCH2* in *B. distachyon* and *OsSPCH1* and *OsSPCH2* in *O. sativa* are partially redundant in their respective species; loss of either paralog leads to a reduction in stomatal production and loss of both eliminates stomata and leads to seedling death (Raissig *et al*., 2016; Wu *et al*., 2019).

Grasses (Poaceae) underwent rapid speciation following the *rho* whole genome duplication (WGD) ∼140-70 million years ago (Lovell *et al*., 2022; Ma *et al*., 2021; Preston *et al*., 2009; Wu *et al*., 2008; Zhang *et al*., 2024). WGDs have been hypothesized to provide the genetic material for novelty because duplication can release genes from selective pressure and enable them to subfunctionalize or acquire new functions (Ohno, 1970). Broadly, plant WGDs contribute to clade survival and environmental robustness on a macroevolutionary timescale (Van de Peer *et al*., 2009). Most plant species return to a diploid state following a WGD (De Smet *et al*., 2013; Panchy *et al*., 2016), yet retain some duplicated genes, and TFs are frequently enriched following plant WGDs (Almeida-Silva and Van de Peer, 2023). Duplicated genes may be retained because they have acquired new functions, they partition ancestral functions, or because gene products are involved in processes where it is critical to maintain stoichiometry (Birchler and Veitia, 2010; Huang *et al*., 2022; Papp *et al*., 2003) TF retention may result when paralogous TFs become expressed in different places, acquire new DNA binding site preferences or become incorporated into new gene regulatory networks (Almeida-Silva and Van de Peer, 2023; Gera *et al*., 2022). The *rho* WGD provided Poaceae with a complex history of duplicate gene retention and losses, and it is estimated that only about one quarter of duplicated genes were retained (Zhang *et al*., 2024). Among the stomatal bHLHs, a second copy of *SPCH*, but not of *MUTE* or *FAMA*, is typically retained in grasses. There are two *SCRM* paralogs derived from a different duplication event than the Brassica-specific duplication that generated *Arabidopsis SCRM* and *SCRM2* (Raissig *et al*., 2016).

In this study, we identify the *rho* WGD as the likely origin of the *SPCH* duplication and provide evidence that both *SPCH* paralogs are under selection. Using loss of function mutations and protein reporters for SPCH1 and SPCH2 in the Pooideae member *B. distachyon*, we show that the two paralogs enable responsiveness to two separate environmental stimuli. To identify the source of paralog-specific regulation, we tested the transcriptional and translational response of each paralog when we altered light and temperature conditions and found that regulation appears to be at the level of protein abundance and persistence in the lineage. Loss of function analyses in the Oryzoideae member, *O. sativa* are consistent with the *SPCH* paralogs mediating similar environmental responses across the grasses.

## Methods

### Synteny analysis

Proteomes, gene locations, and coding domain nucleotide sequences for *Hordeum vulgare* Morex v3, *Triticum aestivum* Chinese Spring v2.1, *Oryza sativa* Nipponbare v7, *Zea Mays* Ref_Gen v4, *Panicum virgatum* v5.1, *Setaria viridis* v4.1, *Sorghum bicolor* v3.1.1 were downloaded from Phytozome.org (accessed 4/11/2025). Orthofinder v2.5 and MCScanX v1.0.0 were used to identify orthologs and blocks of synteny between the grasses. GENESPACE v1.2.3 was used to visualize the blocks of synteny among grass chromosomes and highlight blocks of synteny around *B. distachyon SPEECHLESS* (*SPCH*) paralogs *BdSPCH1* and *bdSPCH2* as a seed for a riparian plot where each species is arranged along the y-axis according to their phylogenetic relationship, accessed from NCBI taxonomy and the regions of synteny. The exclusivity of each of the synteny tracks indicates that the *SPCH* paralogs duplicated before the speciation of these grasses.

### Ka and Ks between paralogs

To calculate the age of the shared grass *SPCH* duplication, we used DIAMOND v2.1.11 blastp program to identify similar proteins within each species using the “–more-sensitive” and “--e 1e-10” flags. Results were filtered to top 5 similar genes for each query gene to improve MCScanX results as recommended in the MCScanX methods paper (Wang *et al*., 2012). MCScanX identified blocks of collinearity and then calculated Ka (nonsynonymous substitutions per site with the possibility of creating a non-synonymous change) and Ks (synonymous substitutions per site of possible synonymous change) between each pair of paralogs in each species. R (v4.2) with the ggplot2 v2.1 package plotted the density for the Ka and Ks results, colored by species and annotating the values for Ka and Ks for the *SPCH1* and *SPCH2* paralogs of each species, respectively.

### Paralog alignment and dN/dS

Coding-domain nucleotide sequences for *SPCH* paralogs from *Hordeum vulgare* Morex v3, *Triticum aestivum* Chinese Spring v2.1, *Oryza sativa* Nipponbare v7, *Zea Mays* Ref_Gen v4, *Panicum virgatum* v5.1, *Setaria viridis* v4.1, and *Sorghum bicolor* v3.1.1 were downloaded from Phytozome. We used TranslatorX to generate a codon-specific nucleotide multi-sequence alignment across all the paralogs (Abascal *et al*., 2010). By creating a nucleotide alignment that preserves codon phase, we could identify the rate of mutations in the nucleotide sequences that lead to synonymous or nonsynonymous changes in the protein. This codon-specific alignment and the phylip gene tree generated by orthofinder were used as inputs into EasyCodeML with the M0 and M2a models(Gao *et al*., 2019; Yang, 2007). For each paralog, we generated Ka/Ks values across each of these genes and plotted using a custom python script in Matplotlib. A negative score across the length of both paralogs indicates that both are still under purifying (negative) selection. Conservation in cis-regulatory elements (CREs) was explored by downloading 600 bp of sequence upstream of the *SPCH1* and *SPCH2* TSSs from Phytozome across 9 grass species (*H. vulgare* r1, *T. aestivum* ChineseSpring v2.1, *O. sativa* Nipponbare v7, *P. hallii* v3.2, *P. virgatum* v5.1, *S. italica* v2.2, *S. viridis* v4.1, *S. bicolor* v5.1, *Z. mays* RefGenV4). Each paralog CRE was aligned with Clustal Omega (Madeira *et al*., 2024). We searched for TF motifs in each CRE using FIMO with the JASPAR Core Plant MEME dataset (Bailey *et al*., 2015; Rauluseviciute *et al*., 2024).

In addition, we aligned amino acid sequences of *SPCH* paralogs across the 11 species from assemblies of the 9 grass species above plus *A. thaliana* (Araport11) and *O. Sativa* (Kitaake v3.1) using CLUSTALW on https://www.genome.jp/tools-bin/clustalw (Thompson *et al*., 1994). We then highlighted the conserved basic helix-loop-helix and SMF (ACT-like) domains in green and yellow, respectively (Seo *et al*., 2022). AtSPCH’s previously described PEST domain is indicated in bold letters, highlighted in pink. We also bolded the putative consensus MAPK phosphorylation sites (P-x-S/T-P) in each paralog (Lewis *et al*., 1998).

### Plant Materials

Multiple *OsSPCH1* and *OsSPCH2* knockout alleles were previously generated in *Oryza sativa L. japonica* cv Zhonghua 11 and described in Wu *et al*., 2019. We used c-*osspch1* and *t-osspch2* (hereafter *osspch1* and *osspch2*) as the representative mutant line for each paralog. CRISPR mutations and a T-DNA insertion were verified by PCR amplification and sequencing (Primers in Table S1). We thank Dr. Suiwen Hou (Lanzhou University) for providing the mutant germplasm. Due to customs constraints and lack of availability of the Zhonghua 11 cultivar in USDA Germplasm Genebank or GRIN-Global database, we were unable to acquire or analyze the Zhonghua 11 wildtype cultivar. We instead analyzed common wildtype cultivars *Oryza sativa L. japonica* cv Nipponbare and *Oryza sativa L. japonica* cv Kitaake. We thank Dr. Julia Bailey-Serres (UC Riverside) and Dr. Zhiyong Wang (Carnegie Institution) for seeds. Multiple CRISPR/Cas9 induced knockout alleles of *BdSPCH1* and *BdSPCH2* were generated previously in *B. distachyon* line Bd21-3 (Raissig *et al*. 2016) and we analyzed a single representative line for each paralog. CRISPR mutations were verified through PCR amplification and sequencing (Primers in Table S1). *B. distachyon* BdSPCH1pro:BdSPCH1-YFP and BdSPCH2pro:BdSPCH2-YFP reporter lines were previously generated in the Bd21-3 line and described in Raissig *et al*. 2016. The Bd21-3 line was used as the *B. distachyon* wildtype.

### Plant growth

Seeds of *B. distachyon* line Bd21-3, and homozygous *bdspch1*, and *bdspch2* mutants were sterilized with 1.2% sodium hypochlorite, 0.1% Tween aqueous solution for 15 minutes then rinsed with sterile water before being placed onto sterile 100mm x 15mm square ½-strength Murashige and Skoog (MS, Caisson Labs) agar plates (1% agar, pH 5.7). We stratified the seeds in darkness at 4°C for 4 days before moving them to a regimen of 8 hours light of 50 µE and 16h dark at 22°C in one of two Percival model CU22L growth chambers (serial numbers: 21160.01.15C and 28704.01.20) for another 2 days for germination. After germination, we moved a third of the plates to a higher light treatment of 300uE and a third to a higher temperature treatment of 32°C in a Percival model CU22L growth chamber. Chamber 21160.01.15C was used for testing response to temperature and chamber 28704.01.20 was used for testing response to light intensity.

*O. sativa* seeds were sterilized with 2.62% sodium hypochlorite in an aqueous solution for 30 minutes before being rinsed in water three times. *O. sativa* seeds were then kept submerged and in darkness at 32°C for 3 days. *O. sativa* was plated 245 x 25mm square on ½-strength Murashige and Skoog (Caisson Labs MS) agar plates (1% agar, pH 5.7) and moved to Percival model CU22L growth chamber (21160.01.15C) with 8 hour light and 16 hour dark cycles with either 50 µE 26°C, 300 µE 26°C, or 50 µE 32°C conditions.

### Stomatal phenotyping

After the third leaf in *B. distachyon* and the second leaf in *O. sativa* fully emerged, we snipped it from the plant and placed it in 7:1 ethanol:acetic acid clearing solution for 1 week. We then mounted leaves on a microscope slide using Hoyer’s solution (7.5 g gum arabic, 5 ml glycerin, 100 g chloral hydrate, 30 ml H2O) and allowed the leaf tissue to clear for one week. DIC images were obtained at 10X for *B. distachyon* leaves and 20X for *O. sativa* leaves magnification on a Leica DMi7 microscope, and three regions per leaf were imaged avoiding the midrib in the fully expanded blade. We then measured the leaf area and cell file length with FIJI-ImageJ and counted the abaxial stomata and stomata cell files with the CellCounter plugin (Schindelin *et al*., 2012).

### RT-qPCR

*B. distachyon* ecotype Bd21-3 seeds were germinated on plates as described above with a daily cycle of 8 hours light of 50µE and 16h dark at 22°C in a Percival model CU22L growth chamber. After 16 days, before the emergence of the third leaf, we moved a third of the plates to either 300 µE 22°C or 50 µE 28°C growth chambers. At each time point of 2 hours, 1 day, 2 days, and 3 days after moving them to the new conditions, we dissected the leaf developmental zone by removing the first two leaves and root, then sectioning 4 mm from shoot apical meristem as previously described (McKown *et al*., 2023; Raissig *et al*., 2016). Tissue was harvested from 6 individuals per replicate with 4 replicates per sample then frozen with liquid nitrogen and stored at -80℃ until all samples were collected. We then ground tissue with a Spex Certiprep Geno Grinder 2000 and extracted RNA from each replicate using the Qiagen RNeasy® Mini kit following the manufacturer’s protocol for total RNA extraction from plant cells, including optional DNase treatment. We quantified the RNA using a Nanodrop (ThermoScientific, NanoDrop 1c) and normalized RNA concentrations across samples. We spiked in 10ng of *A. thaliana* RNA to control for cDNA synthesis variability. To reverse transcribe RNA into single-stranded cDNA, we used the Bio-Rad iScript™ cDNA Synthesis Kit and then quantified the cDNA with Bio-Rad SsoAdvanced™ Universal SYBR® Green Supermix in a CFX96™ Real Time System with primers reported in table S1. Relative expression was computed as 2^−ΔΔ^CT by normalizing Ct values to control gene BdUBC18 and reported as relative to the expression of *BdSPCH1* or *BdSPCH2* in plants remaining at 50 µE 22°C for each the collection time point.

### Fluorescence microscopy of protein reporters

*B. distachyon* BdSPCH1pro:BdSPCH1-YFP and BdSPCH2pro:BdSPCH2-YFP reporter lines were germinated on plates as described above in a daily cycle of 8 hours light (50 µE) at 22°C and 16h dark in a Percival model CU22L growth chamber. After 16 days, before the emergence of the third leaf, we moved a third of the plates to either 300 µE 22°C or 50 µE 28°C chambers. After 3 days, we carefully removed the third leaf, leaving the developmental zone intact by dissecting it out of the surrounding second leaf. We then stained the cell walls with propidium iodide (1:100 of a 1 mg/mL stock) for 7 minutes in darkness then leaf tissue mounted in water. We then imaged the PI and the YFP at 41X on a Leica Stellaris confocal scanning microscope (513nM excitation with 520-570nM emission bandwidth for YFP and 600-650nM emission bandwidth for propidium iodide). We quantified the YFP expression of SPCH1 and SPCH2 protein reporters within the developing leaf using FIJI-ImageJ to identify and record local positions of YFP intensity maxima along the stomata cell file, drawing a 8µm radius circle, and measuring the integral intensity within the circle (Schindelin *et al*., 2012). We measured the length of the zone with fluorescence intensity within each stomata precursor cell file with FIJI-ImageJ. At the base of each stomatal cell file are several cells of uniform shape and size, of which the distal cells will undergo an asymmetric cell division (ACD). The smaller daughter cell of each ACD is fated to become a pair of stomatal guard cells, and at the stages we image leaves, it is possible to see ∼30-50 precursor cells in intermediate stages between the initial ACD and differentiated stoma. We identified the start of the asymmetric cell division (ACD) zone within each cell file as the first YFP+ cell to not be adjacent two other YFP+ cells because ACDs result in larger YFP-cells becoming interspersed in a file. The last YFP+ cell we considered for our measurements was one cell away from the previous YFP+ cell. We ignored the final pre-SCD YFP+ cell in our intensity and length measurements as expression within each cell file typically disappeared several cells before that stage in the lineage before reappearing in that single cell.

## Results

### SPEECHLESS paralogs arose from the *rho* whole genome duplication

In previous studies of grass stomatal development, it was noted that *SPCH* was duplicated, but that MUTE and FAMA, the subfamily Ia bHLHs active at later stages in stomatal development were not (Raissig *et al*., 2016; Wu *et al*., 2019). We were curious whether the two copies of *SPCH* originated from a small-scale duplication of the chromosome around *SPCH*, or whether they were retained after WGD while others were lost. The origin of the *SPCH* duplication would lead us to make different predictions about selective pressures and evolutionary potential of the stomatal lineage gene regulatory networks. Using the regions of *B. distachyon* chromosomes 1 and 3 that contain *BdSPCH1* and *BdSPCH2*, respectively as seeds, we could identify blocks of synteny that contain *SPCH1* and *SPCH2* across the Poaceae family. When visualized in a riparian plot (Fig. 1a), *SPCH* paralogs (*SPCH1* in teal and *SPCH2* in red) appear in distinct, non-overlapping block synteny across multiple species suggesting that the paralogs arose from a large-scale duplication that included neighboring genes. Interestingly, *H. vulgare* (barley) has retained only *SPCH1*.

**Figure 1:**
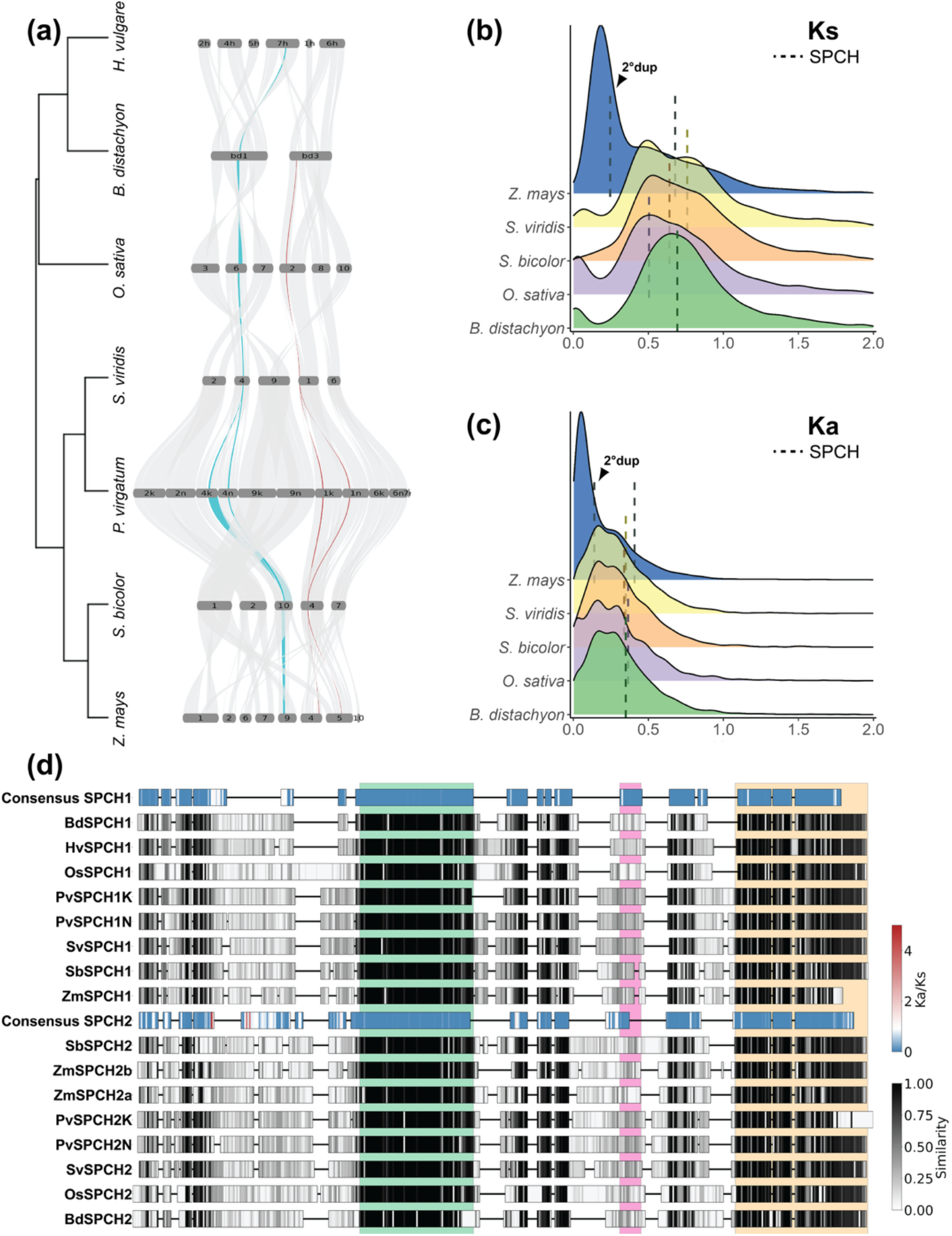
SPCH is retained following the *rho* whole genome duplication in grasses. (a) Riparian plot showing blocks of synteny containing *SPCH1* (teal) and *SPCH2* (red) across multiple grass species. Only chromosomes (Chr) with synteny to *B. distachyon*’s Chr1 and Chr3 are shown and are scaled by the number of genes. (b) Scaled density plot for Ks between paralogs colored by species. Dashed lines represent the Ks value between *SPCH1* and *SPCH2* for the species colored. (c) Scaled density plot of Ka. (d) Amino acid alignment of SPCH1 and SPCH2 proteins across multiple grass species. Consensus sequences for each paralog are colored by Ka/Ks ratio for that amino acid. Paralogs from each species are shaded by similarity across paralogs for that amino acid. Green, pink and yellow underlays represent the bHLH, PEST and SMF (ACT-like) domains, respectively, and correspond to the same color-coded regions in the full amino acid alignment in Fig. S1.

Based on the synteny analysis and distribution of duplicated *SPCH* genes we hypothesized that these paralogs arose from the *rho* WGD. If this were true, then neutral mutation accumulation rates between *SPCH1* and *SPCH2* should be similar to mutation accumulation rates seen between other genes duplicated in this event. From five grass species with high quality genomes, we identified pairs of paralogs within each genome and then calculated the synonymous mutation rate per synonymous site (Ks) and non-synonymous mutation rate per non-synonymous site (Ka) between each paralog pair (Fig. 1b,c). Synonymous mutation rate is often considered to be the rate of neutral evolution (Ohta, 1992). Ks for *SPCH* paralogs in each species is near the center of Ks distribution for other paralogs in each species, indicating that *SPCH* duplicated at the same time as the majority of other retained paralogs (Fig. 1b). This calculated *SPCH* Ks (median, 0.68) is consistent with the Ks calculated for Poaceae-wide paralogs that originated in the *rho* duplication in previous studies (De La Torre *et al*., 2017; Tang *et al*., 2010; Zhang *et al*., 2024). An exception was observed in *Zea mays* (maize) where there is an additional paralog of *SPCH* and the corresponding appearance of a peak at lower Ks values than the other grasses— consistent with the species’ more recent allotetraploid origins (Fig. 1b, blue) (Gaut and Doebley, 1997). Here, our analysis detected both the *pan-*Poaceae duplication that created *SPCH1* and *SPCH2* and a more recent duplication of *SPCH2* (Fig. 1a-c). The Ks values for the more recent duplication of *SPCH2* fall within the peak of *Z. mays* paralogs (Fig. 1b).

If the *SPCH* duplication was a product of a WGD in grasses, then the single copies of *MUTE* and *FAMA* were the result of secondary loss, raising the question of why both *SPCH* copies are retained. Using a metric of selection, the Ka to Ks ratio, we found that non-synonymous mutations are less frequent than synonymous mutations in *SPCH1* and *SPCH2*, indicative of purifying (negative) selection on both genes (Fig. 1c). To identify sites in each gene likely to be the targets of purifying selection, we examined mutation rates separately across *SPCH1* and *SPCH2* paralogs among several Poaceae species, and identified several regions that are conserved and under purifying selection (Fig. 1d). The most conserved of these align with domains shown to be important for function in the single SPCH of *A. thaliana* including the bHLH domain that enables sequence-specific DNA binding (Davies and Bergmann, 2014; Lau *et al*., 2014) and a C-terminal ACT-like domain essential for SPCH activity (Davies and Bergmann, 2014; MacAlister and Bergmann, 2011) that can mediate protein-protein interactions (Figs 1d,S1) (Seo *et al*., 2022). In addition, alignment of the proximal 5’ regulatory regions of *SPCH* paralogs across grass species show that *SPCH1* and *SPCH2* diverged considerably, but that there are blocks of conservation immediately upstream of the TSS of each that contain binding sites of known developmental regulators (Fig. S2).

### Different environmental responsiveness exhibited by *B. distachyon*’s *SPCH* paralogs

*SPCH1* and *SPCH2* acquired molecular differences in both the coding region and within the cis-regulatory regions since their duplication. Because these differences between the paralogs were conserved after speciation across many grasses, we hypothesized that the paralogs acquired functional differences that kept each paralog important and unique. Evidence that both paralogs are under selection is interesting in light of functional studies in *B. distachyon* and *O*.*sativa* showing that SPCH1 and SPCH2 are expressed in and involved in stomatal development, but show unequal redundancy–that is *SPCH2* has a quantitatively larger effect on stomatal production than *SPCH1* and loss of both eliminates stomata completely (Raissig *et al*., 2016; Wu *et al*., 2019). SPCH has a key role integrating environmental conditions into tuning the stomatal lineage in the dicot *A. thaliana*, and so we hypothesized that grass *SPCH* paralogs could sub-functionalize their roles in environmental sensing.

To test the ability of *SPCH1* or *SPCH2* to mediate a change in stomatal production in response to environmental perturbation, we needed to be able to assay plants with only one or the other functional paralog and thus turned to single gene disruption alleles (knockouts) previously characterized in *B. distachyon* (Raissig *et al*., 2016). If either gene mediated the stomatal density (SD, stomata/unit area) response to environmental perturbation, we expected the response to be abrogated in that mutant. In grasses, stomata develop in specific epidermal cell files, and stomata density is a product of the number of cell files in which stomata form and the number of stomata in a given cell file. To measure SD we harvested fully expanded 3rd leaves and imaged the epidermis along the leaf blade, focusing on the developmental zone near the base and avoiding the midrib. For each region imaged, we quantified the number of stomata, the number of stomatal cell files, and the length of each cell file from the initiation to termination in our micrographs to estimate the average distance between stomata in a single cell file (Fig. 2a).

**Figure 2:**
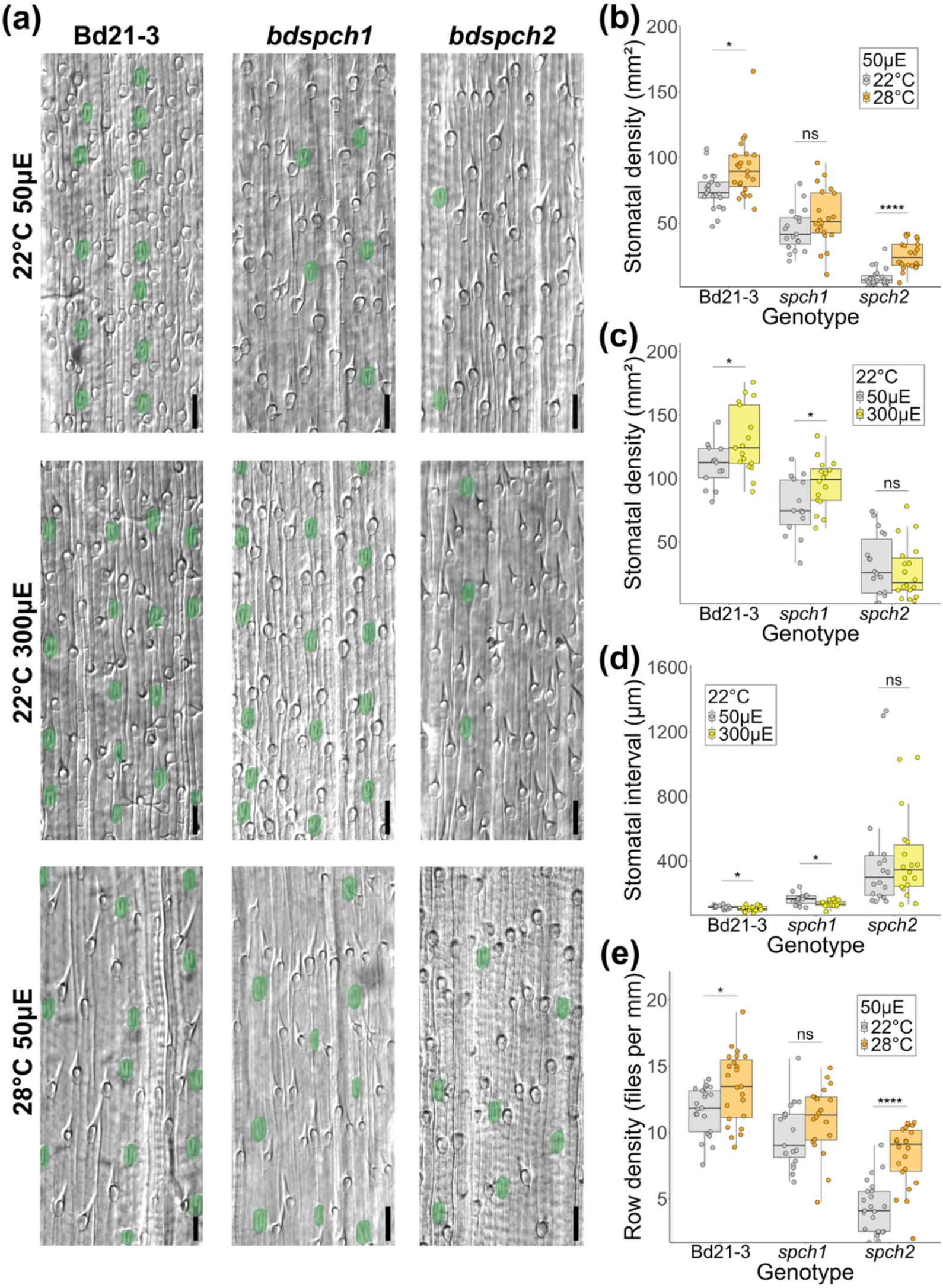
Plants bearing mutations that leave only one *SPCH* paralog functional cannot adjust stomatal density to environmental changes. (a) DIC images of *B. distachyon* abaxial leaf epidermis. Stomata are false-colored green. Images are arranged in rows by environmental conditions and columns by genotype. Scale bar is 50μm. (b) Stomatal density on the abaxial leaf epidermis for plants grown under 22°C or 28°C, 50µE conditions. Plants lacking *BdSPCH1* (*spch1*) are not sensitive to the temperature change. (c) Stomatal density on the abaxial leaf epidermis for plants grown under 50µE or 300µE, 22°C conditions. Plants lacking *BdSPCH2* (*spch2*) are not sensitive to the light intensity change. (d) Average distance (interval) between two stomata in a row. Stomatal spacing decreases with light intensity except in *spch2* plants. (e) The number of epidermal cell files that produce stomata (row density) increases with increased temperature except in *spch1* plants. In (b-e) each point represents a measured developing leaf region (n > 15 regions per genotype and condition). Boxes in panels (b-e) indicate the median (horizontal line) and the interquartile range (box: 25th–75th percentiles). Significance measured by two-way ANOVA with Tukey post-hoc reporting the effect of condition while controlling for leaf identity of each region (* < 0.05, **** < 1e-04).

We established the wildtype Bd21-3 stomatal density response to growth under three different light and temperature regimes (22°C 50 µE, 22°C 300 µE and 28°C 50 µE). The growth conditions generated by changing light intensity and temperature had reproducible effects on stomatal development, with wildtype increasing stomatal density in response to increasing light (50 µE to 300 µE) or temperature (22°C to 28°C) (Fig. 2a-c). In contrast, *bdspch1* mutants failed to show a significant increase in stomatal density in response to warm (28°C) temperature (Fig. 2b) but still responded to an increase in light intensity (Fig. 2c). Conversely, *bdspch2* mutants increased their stomatal density in response to a temperature increase (Fig. 2b) but were insensitive to increased light intensity (Fig. 2c). These reciprocal insensitivities suggest that *B. distachyon SPCH* paralogs have acquired paralog-specific environmental responsiveness. The different insensitivities also allowed us to eliminate the concern that the overall lower number of stomatal in each of the single mutants would render environmental response difficult to generate or to measure. Even in *bdspch2* where overall stomatal production is substantially lower, a response to a change in temperature can be detected.

We found that the cellular mechanism by which SD increased differed between light and temperature change treatments. The distance between stomata in a single cell file decreased significantly when we increased light intensity, but not when we increased temperature (Figs 2d, S3a), except in *bdspch2* where the reverse is true. An increase in the density of stomatal-forming files drove the increase in response to increased temperature (except in insensitive *bdspch1*) but did not change when we increased light intensity (Figs 2e,S3b). Again, the differential behaviors of *bdspch1* and *bdspch2* under these conditions can help pinpoint where in the pathway–from perception of environment to developmental response–the paralogs differ. For example, although *bdspch2* does not decrease spacing in a row in response to a change in light intensity, it can decrease spacing in response to alteration in temperature. Thus, it is likely that *SPCH1* and *SPCH2* can participate in driving similar downstream developmental pathways, and that there is difference in how the upstream environmental cues regulate the paralogs.

### Expression and protein stability of *B. distachyon* paralogs in response to environmental conditions

We then investigated two ways that upstream environmental cues could differentially regulate *BdSPCH1* and *BdSPCH2*. The first hypothesis was *BdSPCH1* and *BdSPCH2* would be transcribed under different conditions, an idea in line with the literature that has shown rapid adaptation through cis-regulatory differences (Fraser *et al*., 2010; Mack *et al*., 2018). Our alignment of 5’ regulatory sequences supported a model that *BdSPCH1* and *BdSPCH2* could be targets of different TFs (Fig. S2), and cell-type specific RNA-seq experiments show that *BdSPCH1* and *BdSPCH2* levels differ across organs and life stages (Hao *et al*., 2021). In Arabidopsis, a stomatal density temperature response is mediated through repression of *SPCH* transcription by the thermosensory TF PIF4 (Lau *et al*., 2018).

To test for a *SPCH* transcriptional response to our alterations of temperature or light, we grew wildtype *B. distachyon* under our standard temperature and light conditions, then moved seedlings to higher temperature or higher light intensity for increasing lengths of time (2 hours to 3 days). *BdSPCH1* and *BdSPCH2* are only expressed in the developing leaf zone (Raissig *et al*., 2016) so we extracted RNA from dissected tissue from this zone. RT-qPCR-based expression profiles showed that RNA abundance of both *BdSPCH* paralogs changed in the same way in response to manipulations of light intensity or temperature at each of the time points we collected following transfer (Fig. 3a). Therefore, we find no evidence that the difference stomata density responses to environmental change observed in the *bdspch1* vs. *bdspch2* is a consequence of different transcriptional regulation of the paralogs.

**Figure 3:**
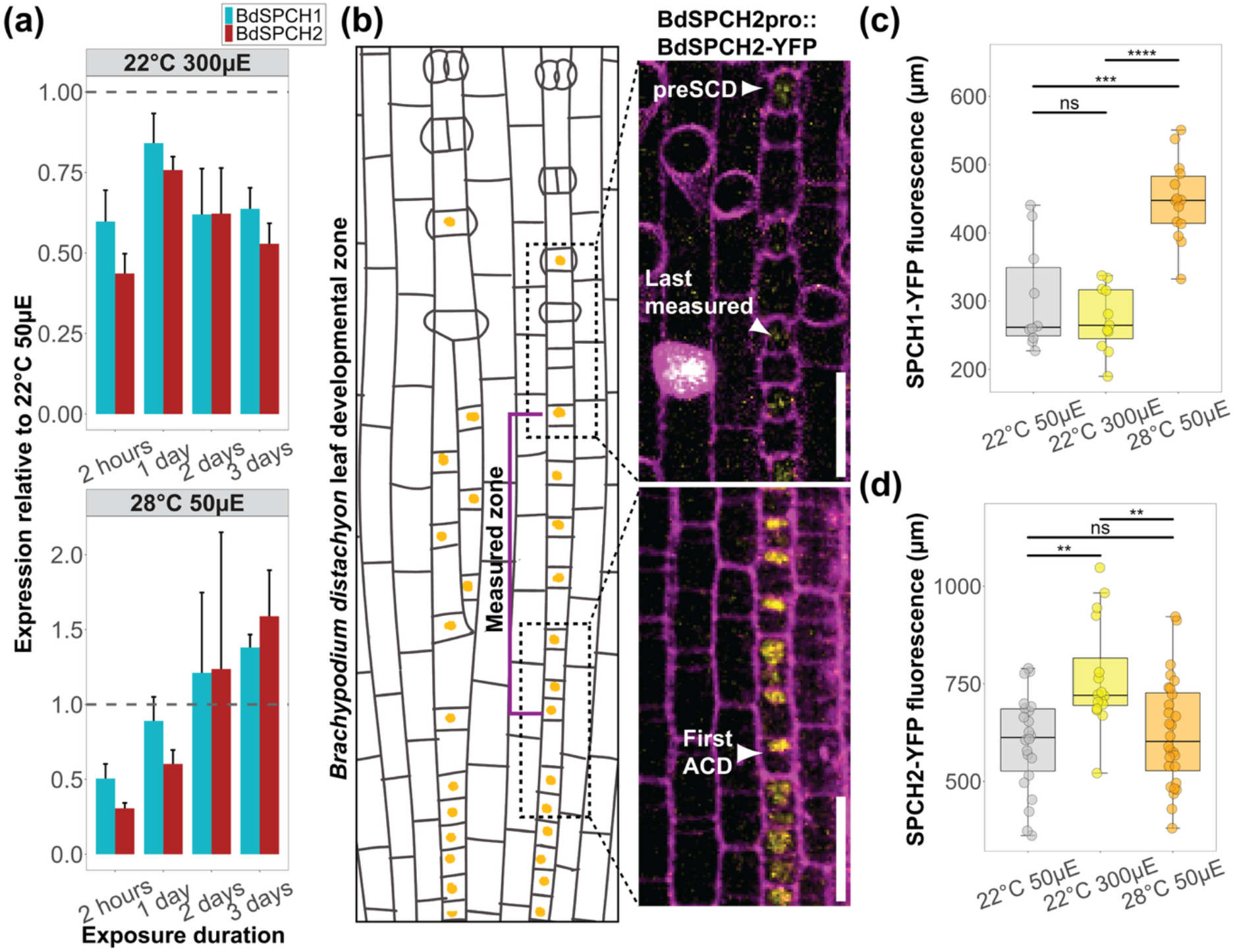
Differential environmental response appears to affect the persistence of SPCH protein, not level of *SPCH* expression. (a) RT-qPCR quantification of *BdSPCH1* and *BdSPCH2* RNA expression after moving *B. distachyon* Bd21-3 plants acclimated to 50µE and 22°C to 300µE light or 28°C temperature for the times specified on the X-axis. Expression quantification was normalized to a control gene and then to the expression of *SPCH1 or SPCH2* isolated from unshifted plants. Bars represent Standard Error (SE). (b) Cartoon and confocal images illustrating *B. distachyon* stomatal lineage development. For this study, BdSPCH2:BdSPCH2-YFP nuclear accumulation (yellow) is quantified from the first asymmetric division (bottom inset, “first ACD”) until the final cell in the contiguous expression domain (top inset, “last measured”). In confocal images, cell outlines are in magenta. Scale bar is 20μm. The cartoon truncates the length of the full developmental zone and ignores hair cells for simplicity. (c,d) Length of detectable BdSPCH1:BdSPCH1-YFP or BdSPCH2:BdSPCH2-YFP fluorescence in “measured zone”. Dots represent cell files across multiple leaves (n > 8 per reporter per condition). Boxes in panels (c) and (d) indicate the median (horizontal line) and the interquartile range (box: 25th–75th percentiles). Significance measured by two-way ANOVA with Tukey post-hoc reporting the effect of condition while controlling for leaf identity of each cell file (** < 0.01, *** < 1e-3, **** < 1e-4).

We therefore tested our second hypothesis, that paralogs are subject to post-translational regulation. Again, there is precedence for this mechanism in the response of Arabidopsis SPCH to environmental factors (Kumari *et al*., 2014; Wang *et al*., 2021) and the grass SPCH1 and SPCH2 alignments show differences in the presence and position of potential phosphorylation sites (Figs 1d, S1) (Raissig *et al*., 2016). We imaged previously generated BdSPCH1pro:SPCH1-YFP and BdSPCH2pro:SPCH2-YFP reporters, and consistent with the prior publications generating these materials (Raissig *et al*., 2016), we detected expression of SPCH1-YFP and SPCH2-YFP in stomatal precursors (Figs 3b,S4a-d). SPCH2-YFP expression appeared in early proliferative cell files and became much stronger in the nuclei of the smaller daughter cells resulting from asymmetric cell divisions (ACDs) in the stomatal cell files. SPCH1-YFP was only detectable after the onset of ACDs. Both SPCH1-YFP and SPCH2-YFP expression decreased as cells matured, but we also noticed a transient burst of nuclear YFP just before the symmetric cell division that creates guard cells (arrow in Fig. S4c,d). This pulse of expression provided a fiducial mark to measure stomatal cell files, but we did not include this cell in our quantifications because fluorescence in the lineage typically disappears a few cells before then (Fig. S4c,d).

Under the same experimental regime as used for assaying transcriptional response, we did identify paralog-specific protein reporter responses to the changes in light intensity and temperature. Specifically, SPCH1-YFP fluorescence persisted longer in stomatal cell files in plants moved to a higher temperature (22°C to 28°C) for 3 days compared to plants moved to a higher light intensity (50 µE to 300 µE) or that remained in the original conditions (Fig. 3c). SPCH2-YFP fluorescence persisted the longest in plants moved to higher light conditions (Fig. 3d). SPCH2-YFP fluorescence could also be detected in proliferating cells at the leaf base but we restricted our quantification to after the first ACD (Fig. 3b) to compare to SPCH1 and focus on confirmed stomatal cell behaviors.

What does extended persistence of BdSPCH reporters mean? In the context of a stomatal cell file, this could be an indication of a prolonged meristematic capability. It could also be a consequence of higher protein accumulation in each cell. By measuring reporter fluorescence intensity per cell, however, we found that the persistence of SPCH1-YFP and SPCH2-YFP was not just due to higher per cell intensity (Fig. S5a,b). In addition, when tracking SPCH2-YFP intensity along a stomatal precursor cell file from base to tip, we found that, under all tested environmental conditions, intensity followed the same low, high, low pattern, with intensity increasing in the middle of the file before trailing off as cells matured (Fig. S5c). We tested whether the length of persistence within the cell file was a consequence of expanding the distance between each stomata precursor. Spacing between the last 10 SPCH1-YFP cells did increase under high temperature conditions, but spacing of the last 10 SPCH2-YFP cells did not significantly increase in any of the environmental perturbations (Fig. S4a,b). Increasing the length of persistence decreases the number of cells between the final pre-SCD cell and second-to-last cell with expression within the cell file (Fig. S4c, distance between yellow bracket and yellow arrowhead).

Overall, reporter expression persistence in stomatal cell files matches the expectations from loss of function mutations as to which SPCH paralog mediates the stomatal density response to our increased light or temperature treatments (e.g., SPCH2 shows longer persistence in higher light and *spch2* mutants were insensitive to increased light). This suggests that environmental regulation specifies not how strongly SPCH paralogs are expressed, but the ability of their proteins to be maintained through development. In Arabidopsis, SPCH protein stability is modulated by MAPK target sites and/or a PEST domain (Davies and Bergmann, 2014; Lampard *et al*., 2008); paralog-specific differences in the presence of these sequences in SPCH1 and SPCH2 (Figs 1d, S1) indicate the potential for this regulatory mechanism to differentially affect paralogs in the grasses.

### Paralog-specific responses to light and temperature are also seen in *O. sativa*

*SPCH* duplicated before the speciation of *B. distachyon* so we were curious whether this subfunctionalization of environmental sensitivity in stomata development was conserved in other grasses that retained both paralogs following the *rho* WGD. *O. sativa* (rice) and *B. distachyon* diverged ∼101-53 MYA and belong to different tribes within the same BOP clade (as distinct from PACMAD containing *Zea mays* and *Sorghum bicolor*) (Christin *et al*., 2014; Zhang *et al*., 2024). Mutations that eliminate *OsSPCH1* or *OsSPCH2* function were previously generated in *O. sativa* (Wu *et al*., 2019) and genomes of several *O. sativa* cultivars with sequence variation around *SPCH* loci are available as well (Fujino *et al*., 2015; Jain *et al*., 2019; Matsumoto *et al*., 2016), thus we tested stomatal development in response to environmental perturbation in loss of function *osspch1* and *osspch2* mutants, and in the *O. sativa* L. *japonica* cultivars Nipponbare and Kitaake.

In *O. sativa*, loss of *OsSPCH2* results in a severe reduction in stomatal density, but loss of *OsSPCH1* was reported to have no discernible stomata density defect (Wu *et al*., 2019). We replicated these results in our chambers and standard growth conditions (Fig. 4a,d,e). We then challenged the *OsSPCH* mutants and Nipponbare and Kitaake cultivars to light and temperature shifts paralleling the shifts performed on *B. distachyon*, but using the reported optimal *O. sativa* growth temperature of 27°C as baseline and 32°C as warm temperature. We compared the responsiveness of *osspch1* and *osspch2* mutants and found that *osspch2* was insensitive to increasing light intensity. Nipponbare, Kitaake and *osspch1* responded by decreasing stomatal density (Fig. 4a-e). The stomata density response to growth in higher light conditions is opposite in direction from that in *B. distachyon* plants, but both *osspch2 and bdspch2 are* insensitive. In response to increased temperature, *osspch2* but not *osspch1* was able to increase stomatal density (Fig. 4a,d,e). Thus, *SPCH1* and *SPCH2* seem to retain similar environmental responsiveness among different species within the BOP grass subfamily. Comparing the behaviors of Kitaake and Nipponbare cultivars in the temperature shift experiment, however, introduced other potential explanations for environmental insensitivity; namely that Kitaake responded similarly to *osspch1* in not responding to increased temperature, leaving the possibility of a cultivar-specific sensitivity to temperature increase in developing leaves (Fig. 4b,c). OsSPCH1 sequences from Kitaake and Nipponbare differ in the presence of a high stringency MAPK target site, providing a potential explanation for the effect, but future experiments would be required to test this (Fig. S1).

**Figure 4:**
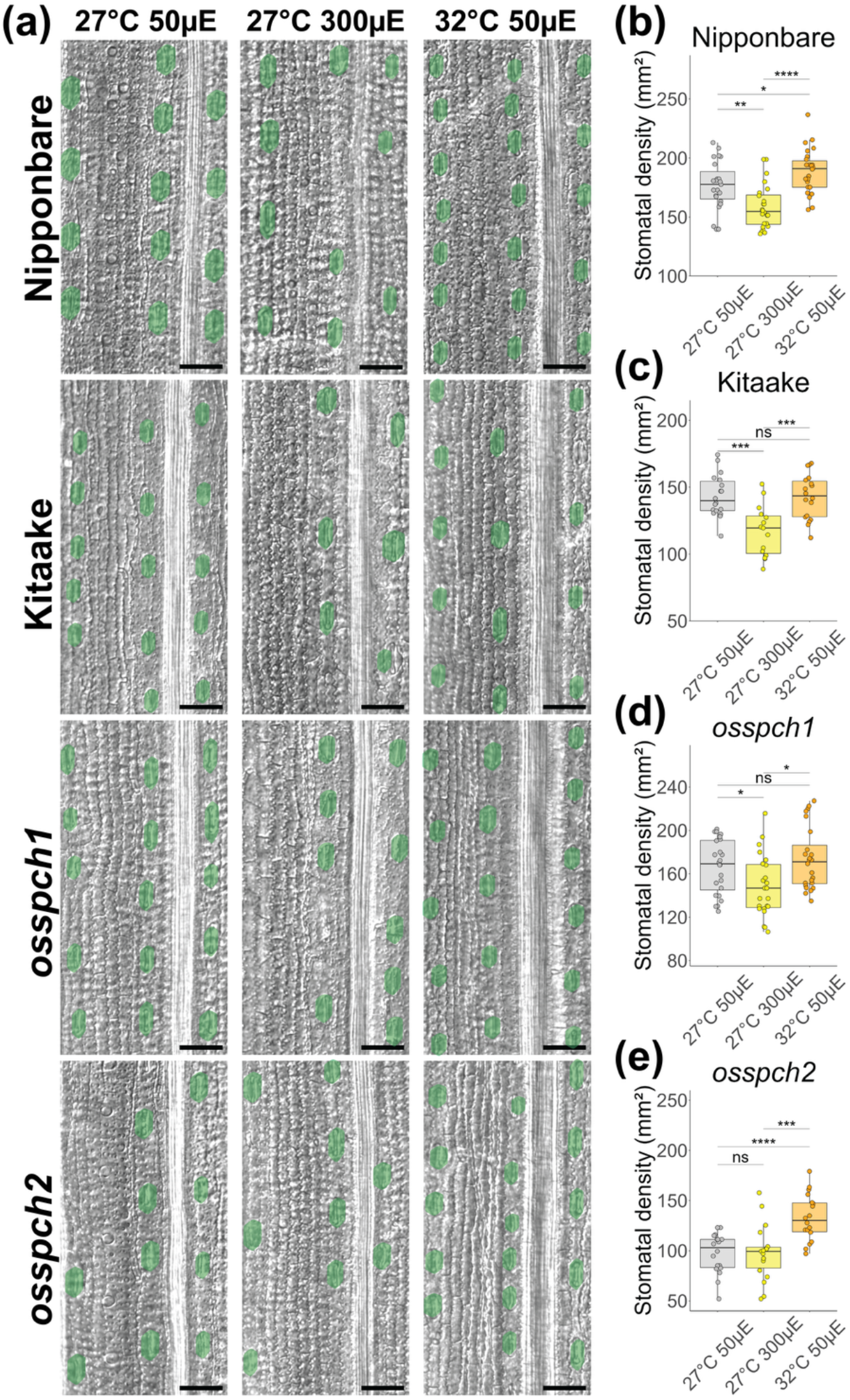
Loss of function *OsSPCH* mutants exhibit the same environmental insensitivities seen with mutations in *B. distachyon SPCH* paralogs. (a) DIC images of *O. sativa* abaxial leaf epidermis. Stomata are false colored green. Images are arranged in columns by environmental conditions and rows by genotype. Scale bar is 50μm. (b-e) Stomatal density on the abaxial leaf epidermis for plants grown under different light or temperature regimes. *OsSPCH1* knockout (KO) and Kitaake plants are not sensitive to the temperature change and *OsSPCH2* KO plants are not sensitive to the light intensity change. Dots represent regions across multiple leaves (Kitaake n > 7, others n > 16). Boxes in panels (b-e) indicate the median (horizontal line) and the interquartile range (box: 25th–75th percentiles). Significance measured by two-way ANOVA with Tukey post-hoc reporting the effect of condition while controlling for leaf identity of each region (* < 0.05,** < 0.01, *** < 1e-3, **** < 1e-4).

Interestingly, when we tracked the cellular mechanism underlying stomatal density adjustments, we found that in *O. sativa* genotypes that respond to changing light intensity, it was due to changing the number of stomatal rows instead of the spacing between stomata within a row (Fig. S6a,b). This differs from the cellular mechanism of response in *B. distachyon* (Figs 2d,S3b). In response to temperature, all *O. sativa* genotypes except Kitaake decrease the distance between stomata within a cell file (Fig. S6a), also different from the observed *B. distachyon* behavior (Fig. 2e,S3a).

## Discussion

The potential for plants to regulate both stomatal aperture and stomatal development enables them to optimize their water use efficiency and photosynthetic capacity across many environments. We show for two representative grasses that stomatal density appears to be tuned in response to environmental cues through varying the number of cell files capable of producing stomata and the number of stomata per file. Changes in stomatal production correlate with differential persistence of SPCH protein accumulation in stomatal files, with SPCH1 and SPCH2 exhibiting unique responses to different environmental conditions. Moreover, we show retention of both copies of the *SPCH* TF originating in the *rho* WGD predating Poaceae speciation may mediate subsets of this response in a conserved manner across BOP clade grasses.

Several mechanisms have been proposed to support the retention of paralogs following WGD in plants including neofunctionalization, subfunctionalization, dosage balance, or neutral variation (Birchler and Yang, 2022; Freeling, 2009; Iohannes and Jackson, 2023). These models can interact and complement each other, for example, dosage balance can maintain redundant paralogs until they acquire subfunctionalization through neutral evolution (Iohannes and Jackson, 2023). While many paralogous TFs were retained from the *rho* WGD, *SPCH* is unique among stomatal lineage bHLH TFs in both copies being retained in most grasses. Each *SPCH* paralog exhibits molecular signatures of purifying selection in the protein coding region (Fig. 1b,d) suggesting an active mechanism maintaining paralogs (Zhang *et al*., 2024). Are any of the proposed general models for paralog retention likely to apply to *SPCH*? A dosage balance model where both *SPCH* paralogs are maintained because reduction in expression would be maladaptive does not fit with previously reported expression data that shows the paralogs have different ratios of expression at different developmental stages (e.g., in embryo vs. along the developmental zone in leaves) (Hao *et al*., 2021; Zhang *et al*., 2022). An extension of this model would suggest that the paralogs would be retained to maintain stoichiometric balance between paralogs and their binding partners (Birchler and Yang, 2022; Kuzmin *et al*., 2020), but this also seems insufficient because the heterodimer partners of SPCH paralogs, SCRM1 and SCRM2, share a more ancestral duplication with other Poales like *Ananas comosus* and are not thought to originate in the *rho* duplication. The phenotypic effects of loss of function mutations in either *SPCH1* or *SPCH2* affect stomatal density rather than other processes, consistent with subfunctionalization rather than neofunctionalization contributing to the retention of these paralogs since the *rho* WGD (Raissig *et al*., 2016; Wu *et al*., 2019).

Functional tests in *B. distachyon BdSPCH1* and *BdSPCH2* mutants grown in different environments support the model of subfunctionalization. Specifically, the paralogs differ in their contributions to the stomatal development response to increasing light intensity or temperature, with *SPCH1* enabling response to the increased light treatment and *SPCH2* to increased temperature. Either paralog could engage the same downstream changes in the density of stomatal cell files or distance between stomata in a cell file, suggesting that it is more likely that the paralogs are subject to different environment-mediated regulation than that they have become enmeshed in different gene regulatory networks to regulate downstream developmental programs.

While we demonstrate subfunctionalization between *BdSPCH* paralogs, we understand a limitation of our approach is that we cannot discern the causal environmental factor in the paralog-specific response to temperature or light intensity. Increasing temperature will increase water vapor pressure and likely increase humidity—additional abiotic factors that have known stomatal density responses (Tricker *et al*., 2012; Devi & Reddy, 2018). In addition, increasing light intensity and temperature both impact photosynthesis within the leaf by either modulating biochemical rates or deactivating the pathway at more extreme conditions (Zhang *et al*., 2002; Scafaro *et al*., 2023). Subsequently, available sugar influences stomata development and decreases stomata density (for example, Gong *et al*., 2021). Regardless of the identity of environmental covariates perturbed by light intensity or temperature changes, plants with mutations inactivating *SPCH1* and those with mutations inactivating *SPCH2* have distinct stomatal responses to the two perturbations, consistent with the SPCH paralogs undergoing subfunctionalization.

Several studies of plant homologs have focused on a transcriptional basis for subfunctionalization following a WGD, thereby expanding our understanding of the evolutionary consequences of duplication (Harikrishnan *et al*., 2015; Li *et al*., 2025). These studies build upon the classic literature tying rapid evolution of regulatory genes to adaptive mutations in cis-regulatory regions (Fraser *et al*., 2010; Mack *et al*., 2018). Indeed, we identified conserved cis-regulatory regions that differ between *SPCH* paralogs across the grasses containing distinct TF motifs (Fig. S2) that could suggest a transcriptional mechanism to explain how paralogs differently affect the phenotype. However, our measured transcriptional differences were insufficient to explain the different dependencies of light and temperature response on the *SPCH* paralogs in *B. distachyon* (Fig. 3a). Neither were average translational intensities sufficient to explain the paralog-specific response to the environment (Fig. S5a,b). Instead, we identified changes in the zone of protein expression that matched the phenotypic behaviors we observed; the zone of protein accumulation within a stomatal file uniquely increased in the reporter of the paralog implicated in mediating a certain environmental response when the plants were exposed to that condition (Fig. 3c,d).

Testing if this subfunctionalization was shared across the BOP clade with *O. sativa* mutants and cultivars revealed a conserved paralog-specific role for *SPCH1* and *SPCH2* in response to temperature and light intensity perturbations, respectively. We note insensitivity to temperature increase seen in *osspch1* was shared with one of two wildtype cultivars we tested–allowing for the possibility that the temperature insensitivity originates in the background cultivar from which *osspch1* was isolated. This explanation, however, would require loss of *OsSPCH2* to have a gain-of-function effect, newly acquiring temperature sensitivity. In *A. thaliana*, different ecotypes display different stomata development responses to changing temperature (Nir *et al*., 2023). The cultivar-specific response underscores the need to test multiple species/cultivars when exploring the environmental regulation of developmental pathways.

Here we present the biological phenomenon of stomatal development responses to environmental change and how it may be mediated through the duplicated *SPCH* genes. The details of how the paralogs can be regulated and how they differ in their regulation is an opportunity for future studies. For example, could this subfunctionalization be adaptive in increasing the species’ environmental tolerance? One approach is to look for correlations between the copy number of *SPCH* paralogs and thermal or latitude ranges in grasses that retained the duplication and the few grasses that did not. This approach of correlating *SPCH* duplications with environmental species range could be expanded across angiosperms–exploring the impact of additional independent duplications of *SPCH* (Danzer *et al*., 2015; McKown *et al*., 2019; Menezes *et al*., 2025) and contributing to our understanding of why TFs involved in environmental responses have increased retention rate following WGD in plants (Panchy *et al*., 2019; Rody *et al*., 2017). For example, *Populus trichocarpa* species have two copies of *SPCH* derived from an independent (from *rho*) duplication, and it appears that alleles of one paralog now predict the ratio of stomata present on the adaxial and abaxial leaf surfaces along an environmental gradient (McKown *et al*., 2019).

Similarly, our study inspires us to consider further functional studies in members of Poaceae that inherited the SPCH duplication but secondarily lost a paralog. *Hordeum vulgare* (barley) and *Triticum aestivum* (wheat) are close relatives of *B. distachyon* in the same subfamily, the “core” Pooideae, but both lost their *SPCH2* paralog and hexaploid wheat now has three copies of *SPCH1*. Are barley and wheat stomatal densities insensitive light intensity changes? If they are light sensitive, could their *SPCH1* paralog have re-acquired post-translational regulation observed in Bd*SPCH2 or OsSPCH2* or have they adopted novel modes of transcriptional regulation?

Finally, the ability to assay reporters of BdSPCH1 and BdSPCH2 protein behavior was essential to consider molecular mechanisms underlying paralog differences. Introducing functional protein reporters of grass SPCH1 and SPCH2 into *A. thaliana spch* mutants could reveal whether the grass paralogs still differ in their capacity to mediate light and temperature stomatal density responses. This heterologous system could also facilitate more detailed investigation of the specific protein domains and residues that mediate environmentally-tuned behavior, with MAPK target sites and a protein-stability determining PEST domain (Lampard *et al*., 2008; Raissig *et al*., 2017; Rogers *et al*., 1986) being strong candidates to test due to their pattern of being conserved among orthologs and distinct between paralogs across grass species (Fig. S1).

Our study contributes a careful examination of the role that paralogs can play in environmental acclimation through non-transcriptional regulation and identifies a novel subfunctionalization in grass paralogs of *SPCH*, a major stomatal development regulator. This contributes to a growing body of knowledge about the mechanisms by which agronomically important plants have or could modify stomatal development in response to changing climates and may contribute to the development of solutions to the challenge of ever-increasing environmental stressors affecting our crop systems.

## Acknowledgements

We thank Zhiyong Wang, Julia Bailey-Serres and Suiwen Hou for rice seeds and Alex Borowsky for advice about rice cultivation. Anastacia Del Rio helped with initial characterization of environmental response and Charlie Hale (Cornell) advised on identifying TF motifs in CRE. Jana Sipkova advised on image processing in ImageJ-FIJI. We thank lab members Katelyn McKown and Gabe Amador for discussions and Alysse Pusey and Genevieve Stier for plant care. JME was supported by funds from the National Institutes of Health (T32GM007276). DCB is an investigator of the Howard Hughes Medical Institute.

## Author contributions

JME contributed to conceptualization, data acquisition, analysis and interpretation and wrote the manuscript in collaboration with DCB. BB contributed to data acquisition. DCB supervised the work.

## Data availability

The data that support the findings of this study, including proteomes, gene locations, and coding domain nucleotide sequences were obtained from Phytozome.org and details of *SPCH* homologues are provided in Table S2. No new sequence data were generated.

