## Supplemental Figures 1-6; tables 1-2 for "SPEECHLESS duplication in grasses expands potential for environmental regulation of stomatal development"

**Fig. S1** Full amino acid alignment of SPCH1 and SPCH2 proteins from selected grass species
**Fig. S2** Predicted paralog-specific transcription factor binding motifs in 5' regulatory sequences
**Fig. S3** Row density and stomatal spacing data for additional genotypes and environments
**Fig. S4** Protein accumulation of BdSPCH paralogs extends along the developmental zone in a paralog-by-condition specific manner
**Fig. S5** SPCH reporter per cell intensity is insufficient to explain paralog-specific environmental response
**Fig. S6** *O. sativa* stomatal density response driven by alternative mechanisms than those used by B. *distachyon*

**Table S1** Primer sequences used for genotyping and expression analysis
**Table S2** Data sources for SPEECHLESS (SPCH) paralog sequences used in this paper

```
                        |         |         |         |         | 1-55
AtSPCH       ---------MQEIIPDFLEECEFVDTSLAGDDLFAILESLEGAGEISPTAASTPK
HvSPCH1      ----MGDALCDQLLVD--VDGEFQ---LPTDAEDLFS-ILETWEDCVNG------
TaSPCH1B     ----MGDALCDQLLVD--VDGEFQ---LPTDAEDLFS-ILETWEDCVHG------
TaSPCH1A     ----MGDALCDQLLVD--VDGEFQ---LPTDAEDLFS-ILETWEDCVNG------
TaSPCH1C     ----MGDALCDQLLVD--IDGEFQ---LPTDAEDLFS-ILETWDDCVNG------
BdSPCH1      ---MGDIALCGQLLLDNVVDGEFQ---LPSDAEDLFS-ILETWEDCVNGSA----
OsSPCH1 (N)  ----MGDALCDQLLLVDSDGGEFIPHHADADADDLFT-ILETWEGCANVVAGGAP
OsSPCH1 (K)  ----MGDALCDQLLLVDSDGGEFIPHHADADADDLFT-ILETWEDCANVVAGGAP
PvSPCH1K     ----MAEALCDQLFSD--VDGELMMMQHPSDADDLLGSILEAWEDCVTGGGGATP
PvSPCH1N     ----MAEALCDQLFSD--VDAGELMMHHASDADDLLG-ILEAWEDCVTGGGCTTP
PhSPCH1      ----MEEALCDQLFSD--VDG-DLMMHHPSDADDLLG-ILEAWEDCVTGGGSTPR
SiSPCH1      ----MAEALCDQLFSD--VDG--ELMHHPSDADDLFG-ILEAWEDCVTGGGSTPR
SvSPCH1      ----MAEALCDQLFSD--VDG--ELMHHPSDADDLFG-ILEAWEDCVTGGGSTPR
SbSPCH1      ----MAEALCDQLFSD--VDG-ELMRQHTSDTDDLLG-ILEAWEDCVTGGGATTP
ZmSPCH1      ----MAEALCDQLFSD--IDG-DLMRQHTSDTDDLLG-ILESWEDCVSGGVATTP
SbSPCH2      ----MADGLCELLVDD---HCELRHRLSG--AEDLFS-IIGTTWKEGTNRAGGDG
ZmSPCH2b     ---MEDGGMCELVIGD---HRNLRHRLSG--AEDLFS-IVGTWEERTNGASGGGG
ZmSPCH2a     ----MGDGQCELVVDN---HCQLRHRLDLSGAEDLFS-VIGTWEERTDDGGAGDG
PvSPCH2K     ---MGDGAMCELLVDD---HSELR--HLS-GAEDLFS-ILETWEECVNGAGGGGG
PvSPCH2N     ---MGDGAMCELLVDD---HSELR--HQSSGAEDLFS-ILETWEECMAGSS----
PhSPCH2      ---MGDGAMCELLVDD---HSELR--HVS-GAEDLFS-ILETWEECMNGGGGGGG
SiSPCH2      ----MADGLCELMVDG---HRDLR--HLS-GAEDLFS-ILETWEECMNGGGGGGG
SvSPCH2      ----MADGLCELMVDG---HRDLR--HLS-GAEDLFS-ILETWEECMNGGGGGGG
OsSPCH2 (N)  MADGGGGGLCELFSDD---HRDIR----HVADADLFR-ILETWEECINGGAGGGG
OsSPCH2 (K)  MADGGGGGLCELFSDD---HRDIR----HVADADLFR-ILETWEECINGGAGGGG
BdSPCH2      MAMGDDGGLCKLFAND---DCEFR-----HVSGDLFS-ILDTWEKCMNGAGEGGG

                  |         |         |         |         |    56-110
AtSPCH       DGTTSSKELVKDQDYENSSPKR--------------------KKQRLETRKEEDE
HvSPCH1      --AAAALSPTAGAGSGGGLLG------------------GSAPGNKGSSKRRARQ
TaSPCH1B     --AAAALSPTAGASSRGELMG------------------GSAPGNK-SSKRRARP
TaSPCH1A     --AAAALSPTAGASSRGGLLG------------------GSAPGNK-GSKRRARQ
TaSPCH1C     --AATALSPTAGASSRGGLLG------------------GSASGNK-GSKRRARQ
BdSPCH1      --ATTALSPTSGASSGALLAPP-----------------DSKLGSKRRMAPQDQE
OsSPCH1 (N)  ATTTTLGSPIAAAACISGVVGGQNHQQLPEPAAAKTVPATNNKRREEEVADRDGD
OsSPCH1 (K)  ATTTTLGSPIAAAACISGVVGGQNHQQLPEPAAAKTVPATNNKRREEEVADRDGD
PvSPCH1K     RSAEPEVPQATDSTARASPPKP----------------------AAGNVRRMGDR
PvSPCH1N     RGGD-EVPQATDSTARASPPKP----------------------AAGNVRRMGDR
PhSPCH1      ---GAKVPQATDAAVATSNPAP----------------------ASGNVRRLGDR
SiSPCH1      G-----AEVSRTANATTPKPPS----------------------AVGSIRRLHDR
SvSPCH1      G-----AEVSRTANATTPKPPS----------------------AVGSIRRLHDR
SbSPCH1      RG---ADVLHQSASDAAATPKA----------------------AAAKRRRQGCR
ZmSPCH1      CG---AEVPQTTAADGAAK----------------------------RRRQGCR
SbSPCH2      AGG-CSSGAMRAYSQSSPAGA-----------------VGARPSVNCRKRTGD--
ZmSPCH2b     GGGGSSAAAVRAYSQGCTAGT-----------------AGAKTAAGTNNSRRR--
ZmSPCH2a     GGGGSSAPAMRAYRQISCSAG-----------------AVAATRLTGKSRRRTGE
PvSPCH2K     SSAAAMPAYSQSSTGGSESAG-----------------AVEGARPTANSRRRAGD
PvSPCH2N     AHAAAMPACSQSSTGGSESAG-----------------AMAGARPTANNRRRAGD
PhSPCH2      GSAAAMPAYSQSSTGGSESAG-----------------AVAGARPTANGRRRAGD
SiSPCH2      GGSSEAMPAYSQSSTGGSDSG-----------------AVAGARPAANSRRRSRD
SvSPCH2      GGSSEAMPAYSQSSTGGSDSG-----------------AVAGARPAANSRRRSRD
OsSPCH2 (N)  GGVSLAGVADQGAAAASTGAGGGA--------------RTTTTTTAANGRRREGR
OsSPCH2 (K)  GGVSLAGVADQGAAAASTGAGGGA--------------RTTTTTTAANGRRREGR
BdSPCH2      GAPCSAAALSPSSVVVVDGAAENN--------------AVVGVRPKPGSRRREAA
```

```
                      |         |         |         |   bHLH 111-165
AtSPCH       EEEDGDGEAEEDNKQDGQQ-----------------------KMSHVTVERNRRK
HvSPCH1      GECDGISHEHKRQKCSPEE-----------------GGGAAPKTAHITVERNRRK
TaSPCH1B     DECDGIAQTQKRQKCSPEE-----------------GGGAAPKTAHITVERNRRK
TaSPCH1A     DECDGIAQAQKRQKCSPEE-----------------GGGAAPKTAHITVERNRRK
TaSPCH1C     DDCDGTAQTHKRQKCSLEE-----------------GGGAAPKTAHITVERNRRK
BdSPCH1      DNDDTAAQAQKRRKCSP----------------------EAPKTAHITVERNRRK
OsSPCH1 (N) GDDDDGSPQKRRKCCSPESS------------TTDVAAATTPKTAHIAVERNRRK
OsSPCH1 (K) GDDDDGSPQKRHKCCSPESS------------TTDVAAATTPKTAHIAVERNRRK
PvSPCH1K     DDDATVPEAPKRQRCSPAMSS------SEAAATSDDGAANNNKTSHITVERNRRK
PvSPCH1N     DDATVPAAPPKRRRCSPAVSS------SEAAATSDDGAASN-KTSHITVERNRRK
PhSPCH1      ED-ATMRAAPKRRRCSPAVSS------SDAAATSDDGAANN-NTSHITVERNRRK
SiSPCH1      DQGDATVPAPKRQRCSPAVSS------EAAAATSEDGAANN-KTSHITVERNRRK
SvSPCH1      DQGDATVPAPKRQRCSPAVSS------EAAAATSEDGAANN-KTSHITVERNRRK
SbSPCH1      EEDGTAVPAPKRQRCSPVSS--------DAAAASEDGAANK--TSHITVERNRRK
ZmSPCH1      DVD-AVPAAPKRQKCSPVS-----------SAASEDGAVNK--TSHITVERNRRK
SbSPCH2      --EEKGSGGGGAPVQKK--------HKGSAVTDDAAADEGEAKMSHITVERNRRK
ZmSPCH2b     --TGDEEKGGSAPAQKK--------HKGSSAVSD---DEGAAKMSHITVERNRRK
ZmSPCH2a     EEEEKGSGGSAPGPAHK--------KHNKAGSAVTDDDEGAPKISHVAVERNRRK
PvSPCH2K     EENGVGRGAPMRKKQKG--------SSTAAAQDAAADEG-AAKMSHIAVERNRRK
PvSPCH2N     EGNGVGRGAPAQKKQKG--------SSTAAAQDVAAADEGGAKMSHIAVERNRRK
PhSPCH2      EEKGVGRGAPAQKKQKG--------SSTAAAAQDAAADEGAVKMSHIAVERNRRK
SiSPCH2      EERGVGRGAPVPKKQKVS-------AAAAITAQDAAADEGAAKMSHIAVERNRRK
SvSPCH2      EERGVGRGAPVPKKQKVS-------AAAAITAQDAAADEGAAKMSHIAVERNRRK
OsSPCH2 (N) DEEKGGGGGGGPPAQKKQKGSSSSSSSPAALAAAVGDGDGAAKMSHITVERNRRK
OsSPCH2 (K) DEEKGGGGGGGPPAQKKQKGSSSSSSSPAALAAAVGDGDGAAKMSHITVERNRRK
BdSPCH2      DEEKGGAPGRK--------------KHKGSTVVDDGSDGAAKMSSHITVERNRRK

                 |         |         |         |         |   166-220
AtSPCH       QMNEHLTVLRSLMPCFYVKRGDQASIIGGVVEYISELQQVLQSLEAKKQRKTYAE
HvSPCH1      QMNEHLTVLRSLMPCFYVKRGDQASIIGGVVDYIKELQQVKQSLEAKKQRKAYTE
TaSPCH1B     QMNEHLTVLRSLMPCFYVKRGDQASIIGGVVDYIKELQQVKQSLEAKKQRKAYTE
TaSPCH1A     QMNEHLTVLRSLMPCFYVKRGDQASIIGGVVDYIKELQQVKQSLEAKKQRKAYTE
TaSPCH1C     QMNEHLTVLRSLMPCFYVKRGDQASIIGGVVDYIKELQQVKQSLEAKKQRKAYTE
BdSPCH1      QMNEHLAALRSLMPCFYVKRGDQASIIGGVVDYIKELQQVKQSLEAKKQRKAYTE
OsSPCH1 (N) QMNENLAVLRSLMPCFYVKRGDQASIIGGVVDYIKELQQVLHSLEAKKQRKVYTD
OsSPCH1 (K) QMNENLAVLRSLMPCFYVKRGDQASIIGGVVDYIKELQQVLHSLEAKKQRKVYTD
PvSPCH1K     QMNEHLAVLRSLMPCFYVKRGDQASIIGGVVDYIKELQQVLQSLEAKKQRKAYTD
PvSPCH1N     QMNEHLAVLRSLMPCFYVKRGDQASIIGGVVDYIKELQQVLQSLEAKKQRKAYTD
PhSPCH1      QMNEHLAVLRSLMPCFYVKRGDQASIIGGVVDYIKELQQVLQSLEAKKQRKAYTD
SiSPCH1      QMNEHLAVLRSLMPCFYVKRGDQASIIGGVVDYIKELQQVLQSLEAKKQRKAYTE
SvSPCH1      QMNQHLAVLRSLMPCFYVKRGDQASIIGGVVDYIKELQQVLQSLEAKKQRKAYTE
SbSPCH1      QMNEHLAVLRSLMPCFYVKRGDQASIIGGVVDYIKELQQVLQSLEAKKQRKAYTE
ZmSPCH1      QMNEHIAVLRSLMPCFYVKRGDQASIIGGVVDYIKELQQVLQSLEAKKQRKAYTE
SbSPCH2      QMNEHLTVLRSLMPCFYVKRGDQASIIGGVVDYIKELQQVLRSLETKKHRKAYAE
ZmSPCH2b     QMNEHLAVLRSLMPCFYVKRGDQASIIGGVVDYIKELQQVLRSLETKKHRKAYAE
ZmSPCH2a     QMNEHLTVLRSLMPCFYVKRGDQASIIGGVVDYIKELQQVLRSLEAKKHRKAYAE
PvSPCH2K     QMNEHLAVLRSLMPCFYIKRGDQASIIGGVVDYIKELQQVLRSLEAKKHRKAYAE
PvSPCH2N     QMNEHLAVLRSLMPCFYIKRGDQASIIGGVVDYIKELQQVLRSLEAKKHRKACAE
PhSPCH2      QMNEHLAVLRSLMPCFYIKRGDQASIIGGVVDYIKELQQVLRSLEAKKHRKAYAE
SiSPCH2      QMNEHLAVLRSLMPCFYVKRGDQASIIGGVVDYIKELQQVLRSLEAKKHRKAYAE
SvSPCH2      QMNEHLAVLRSLMPCFYVKRGDQASIIGGVVDYIKELQQVLRSLEAKKHRKAYAE
OsSPCH2 (N) QMNEHLAVLRSLMPCFYVKRGDQASIIGGVVDYIKELQQVLRSLEAKKNRKAYAD
OsSPCH2 (K) QMNEHLAVLRSLMPCFYVKRGDQASIIGGVVDYIKELQQVLRSLEAKKNRKAYAD
BdSPCH2      QMNENLAVLRTLMPCFYVKRGDQASVIGGVVDYIKELQQVLHSLEAKKHRKVYAV
```

```
                         |         |         |         |     221-275
AtSPCH       VLSPRVVPSP--------RPSPPVLSPRKPPLSPRINHHQIHHHLLLPPISPRTP
HvSPCH1      HVLSPRPPPS--------SYSPRLPLSPLHKSTPPL----SPLLRSTPPLSPRLA
TaSPCH1B     HVLSPRPPPT--------SYSPRLPLSPLHKSTPPL----SPLLRSTPPLSPRLA
TaSPCH1A     HVLSPRPPPS--------SYSPRLPLSPLHKSTPPL----SPLLRSTPPLSPRLA
TaSPCH1C     HVLSPRPPPS--------SYSPRLPLSPLHKSTPPL----SPLLRSTPPLSPRLA
BdSPCH1      QVLSPRPLP---------SPSPRLPLSPLLKSTPPL----SPRLATMS-------
OsSPCH1 (N)  QVLSPRPPAT--------VAASCCSPRPPQLSPRLP----PQLLKSTPPLSPRLA
OsSPCH1 (K)  QVLSPRPPAT--------VAASCCSPRPPPLSPRLP----PQLLKSTPPLSPRLT
PvSPCH1K     QVLSP----------------------RPPLPP---------LKSTPPISPRPS
PvSPCH1N     QVLSPRLPPA--------CCSPRPPLSPRPPLPP---------LKFTPPISPRPS
PhSPCH1      QVLSPRPPPA--------CCSPRPPLSPRPPLPP---------LKSTPPISPRPS
SiSPCH1      QVLSPRPPPP--------SCSPRPPLSPRPPLPP---------LKSTPPISPRPS
SvSPCH1      QVLSPRPPPP--------SCSPRPPLSPRPPLPP---------LKSTPPISPRPS
SbSPCH1      QVLSPR-PPA--------CCSPRPPLSPRPPLPP---------LKSTPPISPRPA
ZmSPCH1      QVLSPR-PPA--------CCSPRPPLSPRPHMLP---------LKSTPPISPRPA
SbSPCH2      QVLSPRPAGG--SVSAASPRPLAAVVKSTPPLSP----------RVAVPISPRTP
ZmSPCH2b     QVLSPRPLP---------------AVKSTPPLSP----------HVAVPMSPRTP
ZmSPCH2a     QVLSPRPSAGGVSTSVSAASPRHLAVKSVAPLSP----------RMAVPISPRTP
PvSPCH2K     QVLSPRPAVP-------AASPR-PLLKSTPPLSP----------RVAVPISPRTP
PvSPCH2N     QVLSPRPAMP-------AASPRPLLIKSTPPLSP----------RVAVPISPRTP
PhSPCH2      QVLSPRPAVS-------AASPR-PLLKSTPPLSP----------RVAVPISPRTP
SiSPCH2      QVLSPRTTVS-------AASPR-PLVKPTPPLSP----------RVAVPISPRTP
SvSPCH2      QVLSPRTTVS-------AASPR-PLVKPTPPLSP----------RVAVPISPRTP
OsSPCH2 (N)  QVLSPRPSPA----------AAALMVKPTPPISPRF----AAAAAAGVPISPRTP
OsSPCH2 (K)  QVLSPRPSPA----------AAALMVKPTPPISPRF----AAAAAAGVPISPRTP
BdSPCH2      EHALSPRPGP---------TPLSPRPLLKPPISP----------RPAVPISPPTP

                  |         |       PEST DOMAIN |         |   276-330
AtSPCH       QPTSPYRAIPPQLPLIPQPPLRSYSSLASCSSLGDPPPYSPASSSSSPSVSSNHE
HvSPCH1      VPISPARTPPTPGSPYKLRPLPP-------PISGSTYVSPAMTPTGYE---PGSS
TaSPCH1B     VPISPARTPPTPGSPYKLRPLPP-------PISGSTYVSPAMTPTGYE---PGSS
TaSPCH1A     VPISPARTPPTPGSPYKLRPLPP-------PISGSTYVSPAMTPTGYD---AASS
TaSPCH1C     VPISPARTPPTPGSPYKLRPLPP-------PISGSTYVSPAMTPTGYD---AASS
BdSPCH1      ----PCRTPPTPGSPYKLIRPLP-------LPPTMSSGSSAYVSPAMT---PTGC
OsSPCH1 (N)  VP-ISPRTPPTPGSPYRLLRLPP-------PPPPASGSNYASPAMTPT---HHET
OsSPCH1 (K)  VP-ISPRTPPTPGSPYRFLRLPP-------PPPPASGSNYASPAMTPT---HHET
PvSPCH1K     VPISPRTPPPGPGSLYKVRMQPP--LPLPLSPPGSAYASPARTPTREPSPASSSS
PvSPCH1N     FPISPRTPPPGPGSPYKVRRQPP--LPLPLSPPGSAYASPARTPTREPSPASSS-
PhSPCH1      VPISPRTPP-GPGSPYKVRRQPP--LPLPLSPPGSAYTSPARTPTREPSPASSS-
SiSPCH1      VLISPRTPP-TPGSPYKLRRQPPPPLPLPLSPPGSAYASPARTPTREPSPAPSY-
SvSPCH1      VLISPRTPP-TPGSPYKLRRQPPSPLPLPLSPPGSAYASPARTPTREPSPAPSY-
SbSPCH1      VPISPRTPPPAPSSPYKPRRQPI--SVPLPPPGSSAYASPAMTPTREP---AAAS
ZmSPCH1      VPISPRTPP-APSSPYKPCR--------LPPPGSSAYASPAMTTTREP---TAAT
SbSPCH2      TPGSPYKPAAAASGAAGTAGSCRLPYPPAAAAYMAAASPAITPTSSSSSYSLDQQ
ZmSPCH2b     TPGSPYKPASGAA--ATTTGSCRLPHRAAAAAAAYIGTPTTSSSSSSYSHDQ---
ZmSPCH2a     TPGSPYK-----AAASGVAGCCRLPLLPPYMLSSSAAAAAASYSHDQQQHYS---
PvSPCH2K     TPGSPYK--PSGGGGGGAASSRPPHPATAACYMMPSPAMTPTTSSSSSSYAHDQQ
PvSPCH2N     TPGSPYKQPSGGGGGSGAAAGSRPPHPAAAAYMMPSPAMTPTTSSSSSSYAHDQQ
PhSPCH2      TPGSPYK----PSGGGGGAGSSRLSHPAAAAYMIPSPAMTPTTSSSSSSYAHDHH
SiSPCH2      TPGSPYKPPPAGGAAAG------SRVPHPAAYMMPSPAMTPATSSSSSSYSHD--
SvSPCH2      TPGSPYKPPPAGGAAAG------SRVPHPAAYKMPSPAMTPATSSSSSSYSHD--
OsSPCH2 (N)  TPGSPYNKHAAAAATARPPHPAAATSSCSVAYSMSPAMTPTSSSSTTTTTTHELS
OsSPCH2 (K)  TPGSPYNKHAAAAATARPPHPAAATSSCSVAYSMSPAMTPTSSSSTTTTTTHELS
BdSPCH2      TPGSPYKPIQRLPHYISPATSSSSTISHAASYDVISPRP----------------
```

```
                      |         |         |         |        331-385
AtSPCH      S-------------------SVINELVANSKS-----------------------
HvSPCH1     --------------YLPSLDAIAAELSVYANRQALQLPPTDLL------------
TaSPCH1B    --------------YLPSLDAIAAELSVYANRQALQLPPTDPL------------
TaSPCH1A    --------------YLPSLDAIAAELSVYANRQALQLPPTDPL------------
TaSPCH1C    --------------YLPSLDAIAAELSVYANRQALQLPPTDPL------------
BdSPCH1     --------------HEPSLEAIAAELSVYAANRQATLLP----------------
OsSPCH1 (N) --------------AAPSLDAIAAELSAYAS--RQALGGGLLL------------
OsSPCH1 (K) --------------AAPSLDAIAAELSAYAS--RQALGGGLLL------------
PvSPCH1K    --------------YLPSLDKIAAELCAYAAGTSKQPQRPALLPAAAAGGGGGAA
PvSPCH1N    --------------YLPSLDKIAAELCAYAAGTSNKPQRPALLPATGGG---GVV
PhSPCH1     --------------YLPSLDKIAAELCAYAAGTNKQQQRQALLPAATSG--GAVV
SiSPCH1     ---------------LPSLDTIAAELCAYAARGTNKQQQAPVALPAAAG---GGG
SvSPCH1     ---------------LPSLDTIAAELCAYAARGTNKQQQAPVALPAAAG---GGG
SbSPCH1     --------------YLPSLDTVAADLCAYAATNK---QLQPALPAAAGG----VV
ZmSPCH1     --------------YLPSLDTIAADLCAYAANKN---KQLQALAAAAGG----DV
SbSPCH2     QQQQHYSMQTTTTTYLPTLDSLVTELAAQAACRPAAAGLN---------------
ZmSPCH2b    --------QRHYSTYLPTLDSLVTELAAQAAACSRPAASGG-------L-----T
ZmSPCH2a    ---------TQTTTYLPTLDSLVTELATQQAACRPAAAAAG-------------L
PvSPCH2K    ----QQLYPTTAQPYLPTLDSLVTELAARAAGG-RPAAAG--------------L
PvSPCH2N    ----QQQHYPTTQLYLPTLDSLVTELAARAAGG-RPAAAG--------------L
PhSPCH2     ----QQHYP-TTQPYLPTLDSLVTELAAQAAGG-RPAAAG--------------L
SiSPCH2     ----QQHYPATSQPYLPTLDSLVTELAAQAAGAGRPAAAG--------------L
SvSPCH2     ----QQHYPATSQPYLPTLDSLVTELAAQAAGAGRPAAAG--------------L
OsSPCH2 (N) ----------PAPAFLPILDSLVTELAARGGASCRPLVIPSSAAAIAGI-----V
OsSPCH2 (K) ----------PAPAFLPILDSLVTELAARGGASCRPLVIPSSAAAIAGI-----V
BdSPCH2     --------------YLPTLDSIITELAAQAAARAPGALGGGGGGAGLNL-----L

             SMF/ACT DOMAIN|         |         |         |   386-440
AtSPCH      ALADVEVKFSGANVLLKTVSHKI-PGQVMKIIAALED--LALEILQVNINTVDET
HvSPCH1     --PDVRVEFAGANLVLKTVSHRA-PGQAVKIIAALESRAPALEILHAKISTIDDT
TaSPCH1B    --PDVRVEFAGANLVLKTVSHRA-PGQVVKIIAALESRAPALEILHAKISTIDDT
TaSPCH1A    --PDVRVEFAGANLVLKTVSHRA-PGQAVKIIAALESRAPALEILHAKISTIDDT
TaSPCH1C    --PDVRVEFAGANLVLKTVSHRA-PGQAVKIIAALESRAPALEILHAKISTIDDT
BdSPCH1     ---DVRVEFRGANLVLKTVSPRA-PGQAVKIVAALEG--RALEILHAKISTVDDT
OsSPCH1 (N) --PDVKVEFAGANLVLKTVSQRS-PGQAVKIIAALEG--RSLEILHAKISTVDDT
OsSPCH1 (K) --PDVKVEFAGANLVLKTVSQRS-PGQAVKIIAALEG--RSLEILHAKISTVDDT
PvSPCH1K    LLPDVRVEFSGANLVVKTVSHRA-PGQAVKVIAALEG--RSLEILDAKISTVDDT
PvSPCH1N    LLPDVRVEFSGANLVVKTVSHRA-PGQAVKVIAALEG--RSLEILDAKISTVDDT
PhSPCH1     LLPDVRVEFSGANLVVKTVSHRE-PGQAVKVIAALEG--RSLEILDAKISTVDDT
SiSPCH1     LLPDVKVEFAGANLVVKTVSHRA-PGQAVKIIAALEG--RSLDILDAKISTVDDT
SvSPCH1     LLPDVKVEFAGANLVVKTVSHRA-PGQAVKIIAALEG--RSLDILDAKISTVDDT
SbSPCH1     VLPDVKVEFSGANLVVKTVSHRA-PGQTVKVIAALEG--RSLEILDAKINTVNDT
ZmSPCH1     VLPDVKVEFSGANLVVKTVSHRA-PGQTVKVIAALEG--RSLEILDAKINTINDT
SbSPCH2     -LPDVKVEFAGPNLVLKTVSHRS-PGQALKIIAALES--LSLEILHVSVSTVDDT
ZmSPCH2b    RLPDVKVEFAGPNLVLKTVSHRS-PGQALKIIAALES--LPLEILHVSVSTVDDT
ZmSPCH2a    ALPDVKVEFAGPNLVLKTVSHRA-PGQALKIIAALES--LSLQILHVSVSAVDDT
PvSPCH2K    TLPDVKVEFAGPNLVLKTVSRRA-PGQALKIIAALES--LSLEILHVSVSTLDDT
PvSPCH2N    TLPDVKVEFAGPNLVLKTVSRRA-PWQALKIIAALES--LSLEILHVSVSTLDDT
PhSPCH2     ALPDVRVEFAGPNLVLKTVSHRA-PGQALKIIAALES--LSLEILHVSVSTLDDT
SiSPCH2     ALPDVRVEFAGPNLVLKTVSHRA-PGQALKIIAALES--LSLEILHVSVSTVDDT
SvSPCH2     ALPDVRVEFAGPNLVLKTVSHRA-PGQALKIIAALES--LSLEILHVSVSTVDDT
OsSPCH2 (N) GVPDVRVEFAGPNLVLKTVSHRA-PGQALKIIAALES--LSLEILHVSICTVDDA
OsSPCH2 (K) GVPDVRVEFAGPNLVLKTVSHRA-PGQALKIIAALES--LSLEILHVSICTVDDA
BdSPCH2     LLPDVKVEFAGPNLVLKTTSHRARPGQVLRIIAALES--LSLEILHVSISTVDDT
```

```
                      |         |      441-473
AtSPCH        MLNSFTIKIGIECQLSAEELAQQIQQTFC----
HvSPCH1       DVNAFTVKIGIECSLSAEELVQEIQQTFS----
TaSPCH1B      DVNAFTVKIGIECSLSAEELVQEIQQTFS----
TaSPCH1A      DVNAFTVKIGIECSLSAEELVQEIQQTFS----
TaSPCH1C      DVNAFTVKIGIECSLSAEELVQEIQQTFS----
BdSPCH1       DVNAFTVKIGIECELSAEELVQEIQQTFS----
OsSPCH1 (N)   AVNSFTVKIGIECELSAEELVQVIQQTFT----
OsSPCH1 (K)   AVNSFTVKIGIECELSAEELVQVIQQTFT----
PvSPCH1K      AVNSFTIKIGIECELSAEELVQEIQQAFS----
PvSPCH1N      AVNSFTIKIGIECELSAEELVQEIQQAFS----
PhSPCH1       AVNSFTIKIGIECELSAEELVQEIQQAFS----
SiSPCH1       AVNSFTIKIGIECELSAEELVQEIQQAFS----
SvSPCH1       AVNSFTIKIGIECELSAEELVQEIQQAFS----
SbSPCH1       AVNSYTIKIGIECELSAEELVQEIQQAFSS---
ZmSPCH1       AVNSYTIKVLLSSS-------------------
SbSPCH2       MVHSFTIKIGIECELSAEELVQEIQQTLL----
ZmSPCH2b      MVHSFTIKIGIECELSAEELVQEIQQTLL----
ZmSPCH2a      MLHSFTIKIGIECELSAEELVQEIQQTLL----
PvSPCH2K      MVHSFTIKVYIDSLFITWCLIVHNNFLIIHVVY
PvSPCH2N      MVHSFTIKIGIECELSAEELVQEIQQTFL----
PhSPCH2       MVHSFTIKIGIECELSAEELVHEIQQTFL----
SiSPCH2       MVHSFTIKIGIECELSAEELVHEIQQTLL----
SvSPCH2       MVHSFTIKIGIECELSAEELVHEIQQTLL----
OsSPCH2 (N)   TVLSFTIKIGIECELSAEELVQEIQQTFL----
OsSPCH2 (K)   TVLSFTIKIGIECELSAEELVQEIQQTFL----
BdSPCH2       MVHSFTIKIGIECELSAEELVQEIRQTFL----
```

**Fig. S1 Full amino acid alignment of SPCH1 and SPCH2 proteins from selected plant species.** Two *Oryza sativa* L. *japonica* cultivars are included and differentiated by (K) for Kitaake and (N) for Nipponbare, SPCH2 proteins are denoted in lighter grey. Basic-helix-loop-helix (bHLH) is highlighted in green. PEST domain in *Arabidopsis thaliana* annotated with pink highlight and bold text. Putative consensus MAPK phosphorylation sites (P-x-S/T-P) are marked by bold text. SMF (also known as ACT-like) domain is highlighted in yellow. Alignment was performed with CLUSTALW on https://www.genome.jp/tools-bin/clustalw on genes listed in Table S2.

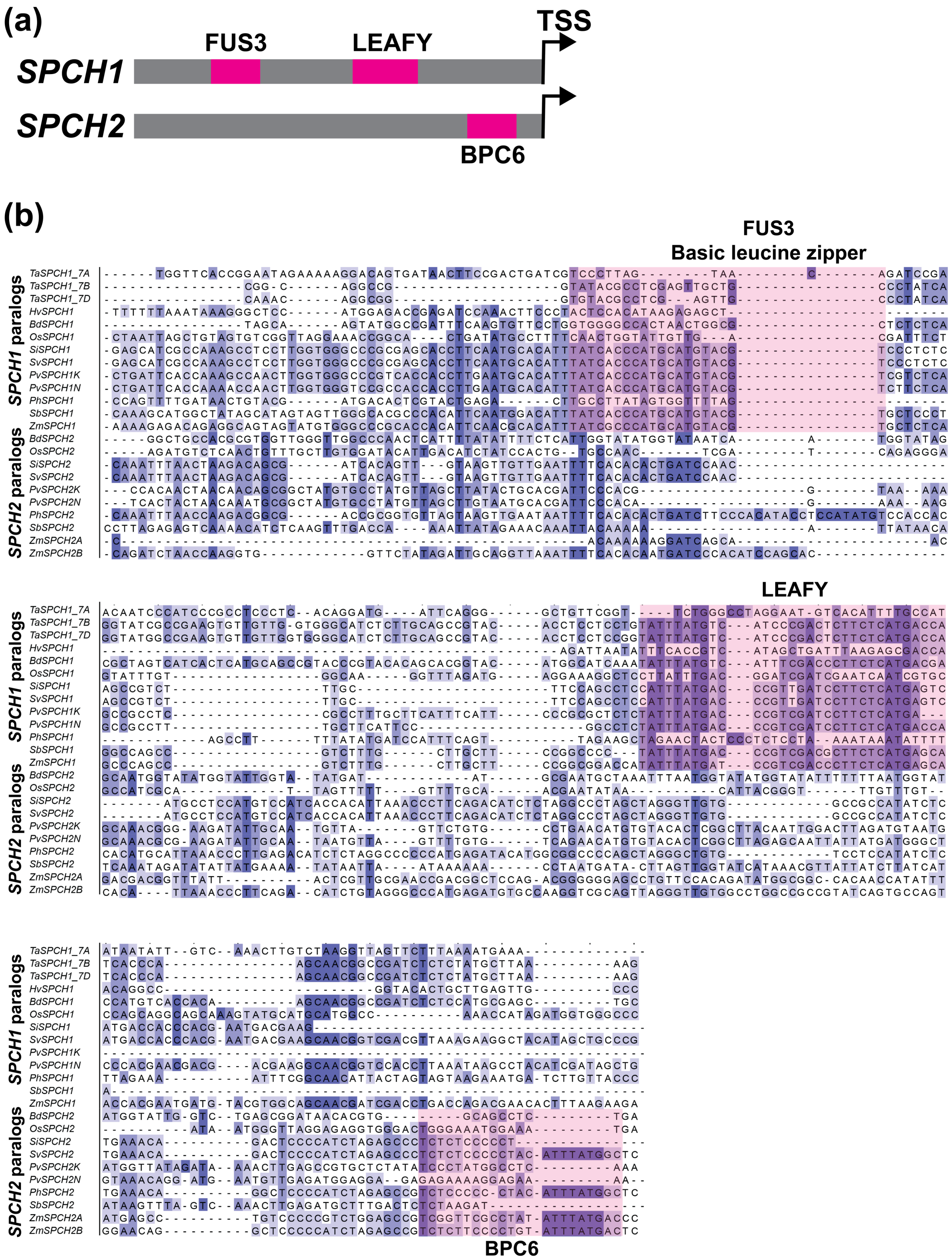


**Fig. S2 Predicted paralog-specific transcription factor binding motifs in 5' regulatory sequences.** (a) Overview of alignment features and orientation upstream of the transcription start site for *SPCH* paralogs. (b) Section of alignment of 600 bp upstream of the *SPCH* transcription start site for *SPCH* paralogs in *B. distachyon*, *H. vulgare*, *T. aestivum*, *O. sativa*, *P. hallii*, *P. virgatum*, *S. italica*, *S. viridis*, *S. bicolor*, and *Z. mays*. Alignment was performed with Clustal Omega and conserved regions containing TF motifs identified by FIMO with the JASPAR core plant motif dataset are highlighted in pink.

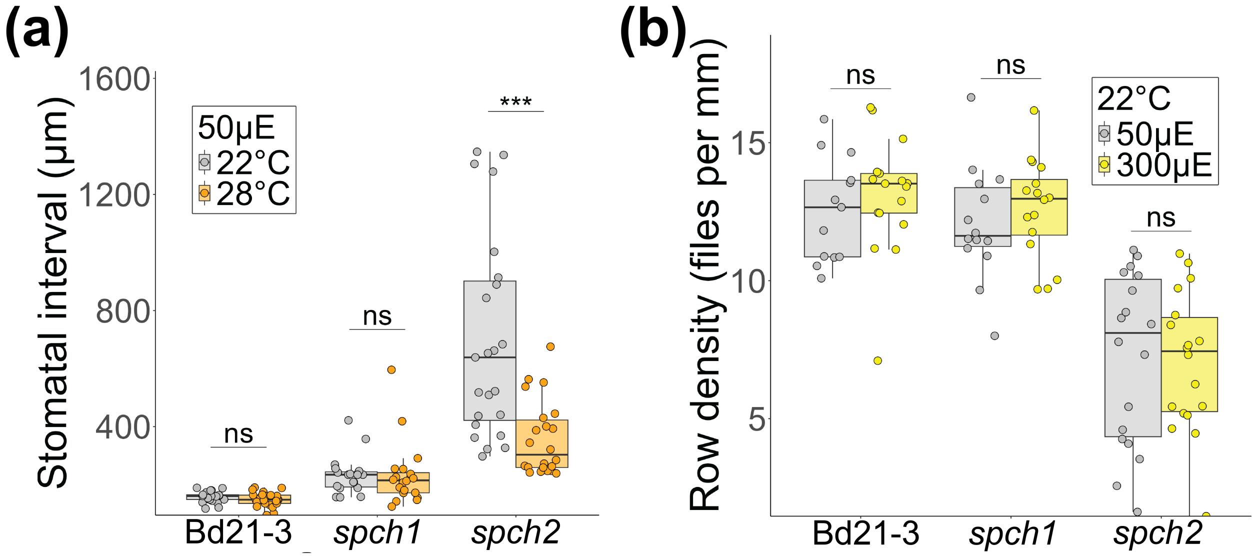


**Fig. S3 Row density and stomatal spacing data for additional genotypes and environments.** (a) Average space (interval) between two stomata in a row in *B. distachyon* plants grown at 22°C or 28°C, 50µE conditions. Stomata spacing does not decrease with temperature except in *bdspch2* plants. (b) Row density of stomata cell files in plants grown at 50µE or 300µE, 22°C conditions. Row density does not increase with light intensity. Dots represent regions across multiple leaves (n > 15 per genotype and condition). Boxes in panels (a) and (b) indicate the median (horizontal line) and the interquartile range (box: 25th–75th percentiles). Outliers are plotted as individual points. Two-way ANOVA with Tukey post-hoc reporting the effect of condition while controlling for leaf identity of each region (*** < 1e-3).

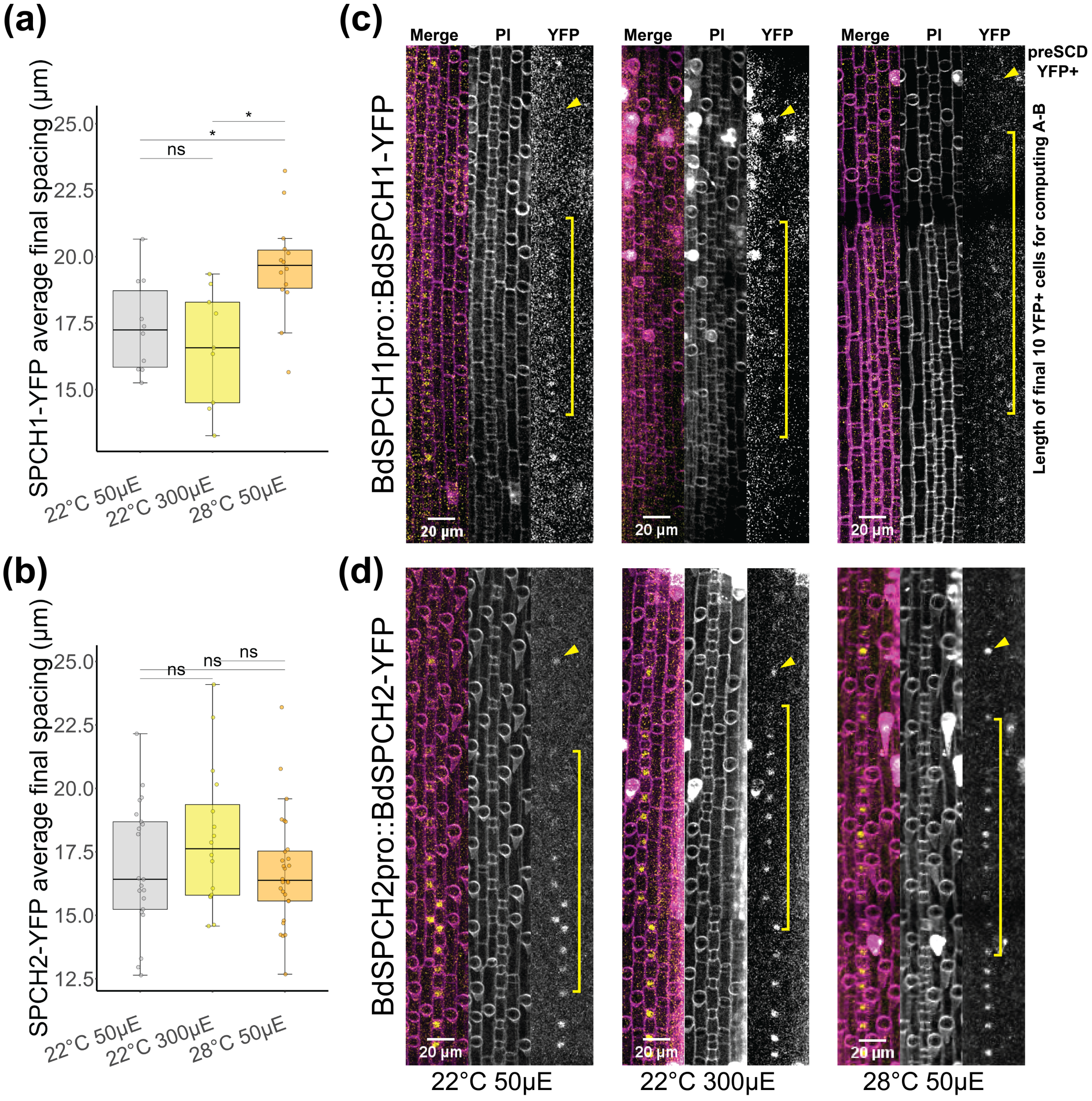


**Fig. S4 Protein accumulation of BdSPCH paralogs extends along the developmental zone in a paralog-by-condition specific manner.** Average space between the ten YFP-positive cells before the final pre-SCD cell in (a) BdSPCH1:BdSPCH1-YFP or (b) BdSPCH2:BdSPCH2-YFP. Dots represent cell file averages across multiple *B. distachyon* leaves (n > 8 per genotype and condition). Boxes in panels (a) and (b) indicate the median (horizontal line) and the interquartile range (box: 25th–75th percentiles). Two-way ANOVA with Tukey post-hoc reporting the effect of condition while controlling for leaf identity of each cell file (* < 0.05). (c,d) Representative epidermal micrographs illustrating distribution of YFP fluorescence in each cell file before final pre-SCD expression. Yellow brackets indicate examples of 10 measured cells and yellow arrowheads indicate pre-SCD cell.

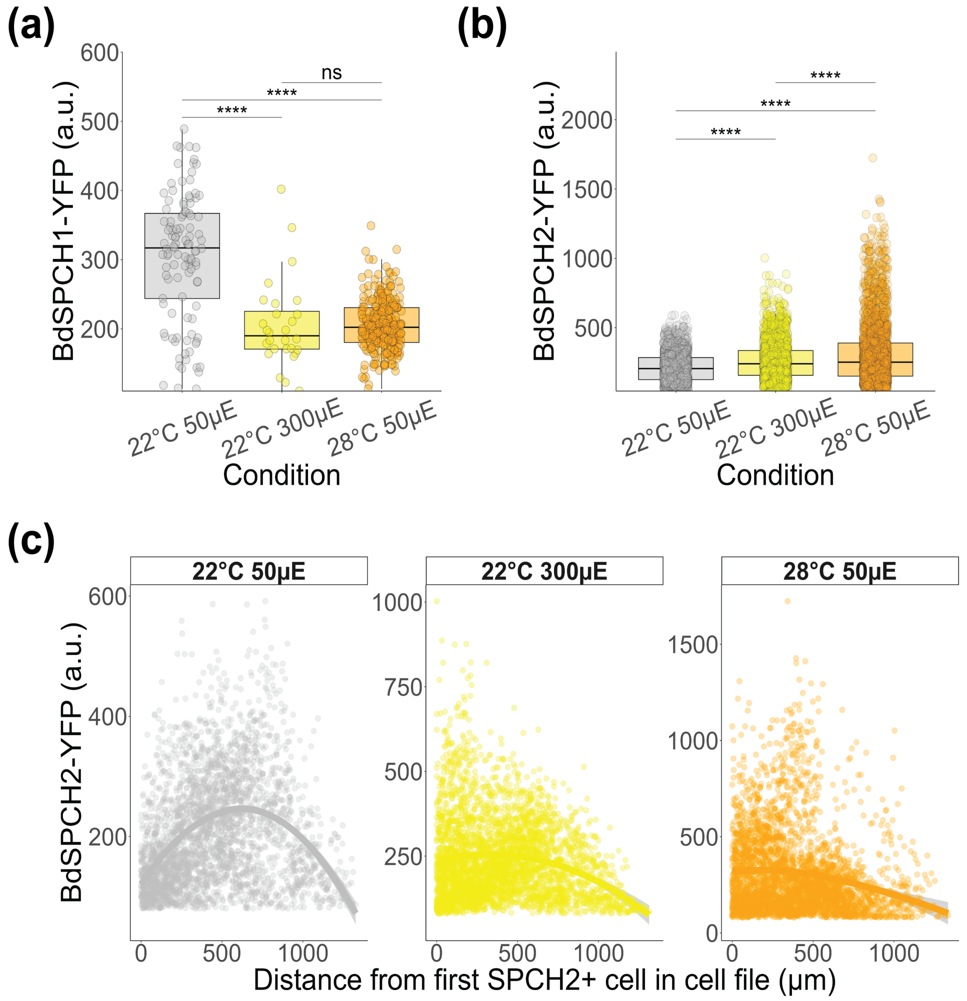


**Fig. S5 SPCH reporter per cell intensity is insufficient to explain paralog-specific environmental response.** Intensity of reporter fluorescence on a per cell basis under multiple conditions from (a) BdSPCH1:BdSPCH1-YFP or (b) BdSPCH2:BdSPCH2-YFP Two-way ANOVA with Tukey post-hoc (**** < 1-e4) reporting the effect of condition while controlling for *B. distachyon* leaf identity (n > 8 per genotype and condition) of each cell (n > 100). Boxes in panels (a) and (b) indicate the median (horizontal line) and the interquartile range (box: 25th–75th percentiles). (c) Cell fluorescence intensity of BdSPCH2:BdSPCH2-YFP by distance from the first fluorescent cell within that cell file summed across all cell files for each condition. Smoothed conditional means line (LOESS regression) shows shared trend across conditions of increasing and then decreasing fluorescence intensity.

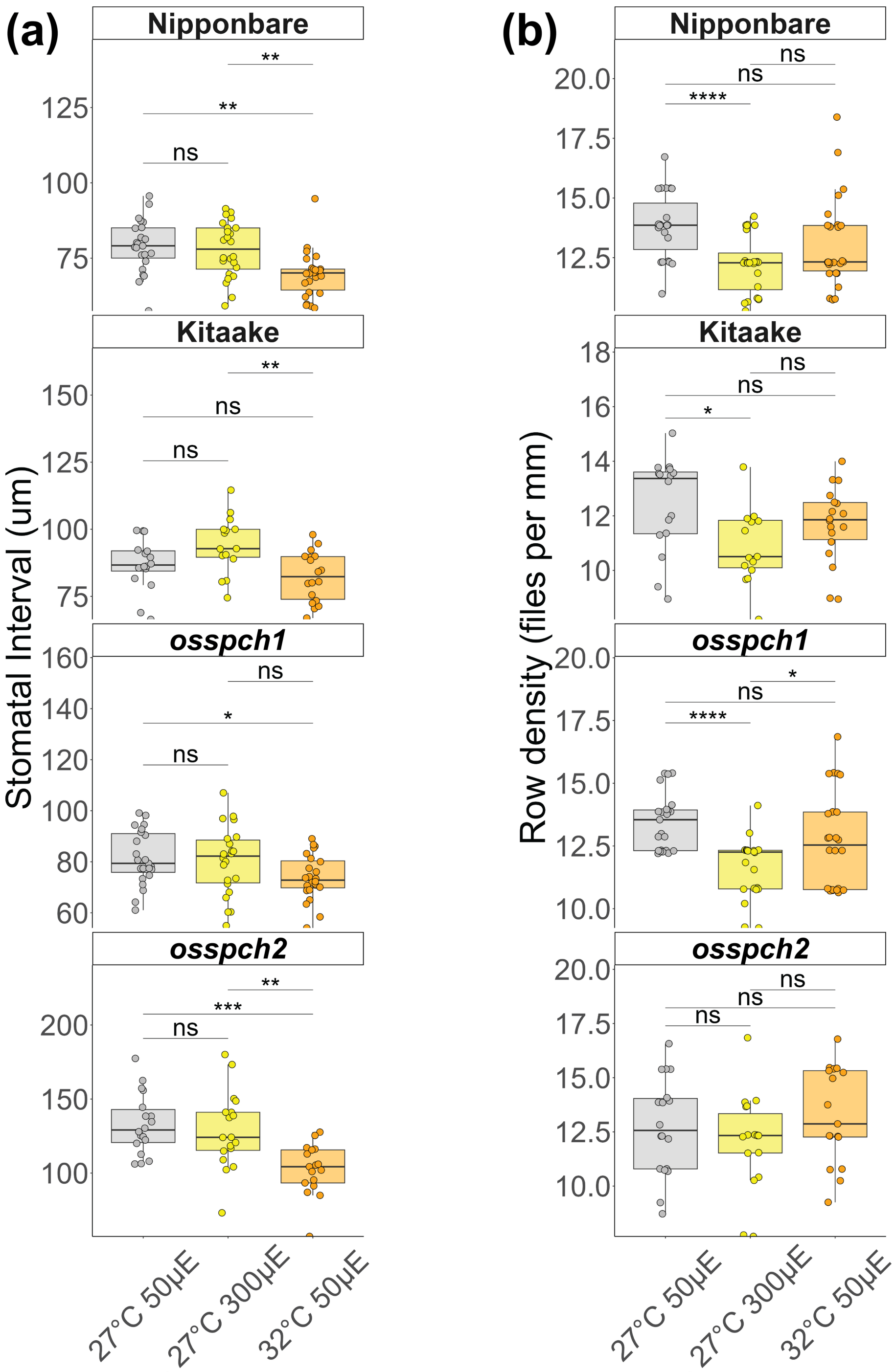


**Fig. S6 *O. sativa* stomatal density response driven by alternative mechanisms than those used by B. *distachyon.*** (a) Average space (interval) between two stomata in a row across *O.sativa* genotypes and conditions. (b) Row density of stomata cell files across *O.sativa* genotypes and conditions. Dots represent regions across multiple leaves (Kitaake n > 7, others n > 16 per condition). Boxes in panels (a) and (b) indicate the median (horizontal line) and the interquartile range (box: 25th–75th percentiles). Two-way ANOVA with Tukey post-hoc reporting the effect of condition while controlling for leaf identity of each region (* < 0.05, ** < 0.01, *** < 1e-3, **** < 1e-4).

**Table S1 Primer sequences used for genotyping and expression analysis**

| **Primer Name** | **Primer Sequence** | **Purpose** |
|---|---|---|
| bdspch1-F | GCACGTACACCACTGATCAT | *bdspch1* genotyping |
| bdspch1-R | ACATAGAAGCACGGCATGAG | *bdspch1* genotyping |
| bdspch2-F | ACACATGCACAGAACGACCT | *bdspch2* genotyping |
| bdspch2-R | AACTCCACCTTCACGTCAGG | *bdspch2* genotyping |
| osspch1-F | CGACCTCTTCACCATCCTCG | *osspch1-1-c* genotyping |
| osspch1-R | GGCGAGGTTCTCGTTCATCT | *osspch1-1-c* genotyping |
| osspch2-F | AACGGTCTACTCTCTCTCTCTC | *osspch2-t* genotyping |
| osspch2-R | GCACTGTAGTACTCTTGGTCAC | *osspch2-t* genotyping |
| BdSPCH1qpcr-F | CACCGTCAAGATTGGAATCGAGTG | qPCR primer for *BdSPCH1* |
| BdSPCH1qpcr-R | TCACGAGAACGTTTGCTGAATCTC | qPCR primer for *BdSPCH1* |
| BdSPCH2qpcr-F | TAGCACCGTCGACGACACTATG | qPCR primer for *BdSPCH2* |
| BdSPCH2qpcr-F | TTCGGCGCTAAGCTCACATTCG | qPCR primer for *BdSPCH2* |
| BdUBC18qpcr-F | GTCACCCGCAATGACTGTAAGTTC | qPCR primer for *BdUBC18* |
| BdUBC18qpcr-R | TTGTCTTGCGGACGTTGCTTTG | qPCR primer for *BdUBC18* |
| AtUBC18qpcr-F | CAGTCTGTGTGTAGAGCTATCA | qPCR primer for *AtUBC18* |
| AtUBC18qpcr-R | ATCTTAGAAGATTCCCTGAGTC | qPCR primer for *AtUBC18* |

**Table S2 Data sources for *SPEECHLESS (SPCH)* paralog sequences used in this paper**

| **Species** | **Gene label** | **Gene Id** | **Genome version (accessed from phytozome.org)** |
|---|---|---|---|
| *Brachypodium distachyon* | SPCH1 | Bradi1g38650 | v3.2 |
| *Brachypodium distachyon* | SPCH2 | Bradi3g09670 | v3.2 |
| *Hordeum vulgare* | SPCH1 | HORVU7Hr1G079250 | r1 |
| *Oryza sativa* Nipponbare | SPCH1 | LOC_Os06g33450 | v7 |
| *Oryza sativa* Nipponbare | SPCH2 | LOC_Os02g15760 | v7 |
| *Oryza sativa* Kitaake | SPCH1 | OsKitaake06g172500 | v3.1 |
| *Oryza sativa* Kitaake | SPCH2 | OsKitaake02g112600 | v3.1 |
| *Panicum hallii* | SPCH1 | Pahal.4G131400 | v3.2 |
| *Panicum hallii* | SPCH2 | Pahal.1G118900 | v3.2 |
| *Panicum virgatum* | SPCH1K | Pavir.4KG254300 | v5.1 |
| *Panicum virgatum* | SPCH1N | Pavir.4NG174200 | v5.1 |
| *Panicum virgatum* | SPCH2K | Pavir.1KG252700 | v5.1 |
| *Panicum virgatum* | SPCH2N | Pavir.1NG142600 | v5.1 |
| *Setaria italica* | SPCH1 | Seita.4G167700 | v2.2 |
| *Setaria italica* | SPCH2 | Seita.1G013200 | v2.2 |
| *Setaria viridis* | SPCH1 | Sevir.4G135400 | v4.1 |
| *Setaria viridis* | SPCH2 | Sevir.1G013100 | v4.1 |
| *Sorghum bicolor* | SPCH1 | Sobic.010G152700 | v5.1 |
| *Sorghum bicolor* | SPCH2 | Sobic.004G115000 | v5.1 |
| *Triticum aestivum* | SPCH1A | TraesCS7A03G0857600 | ChineseSpring v2.1 |

| | | | |
|---|---|---|---|
| *Triticum aestivum* | SPCH1B | TraesCS7B03G0653200 | ChineseSpring v2.1 |
| *Triticum aestivum* | SPCH1C | TraesCS7D03G0780300 | ChineseSpring v2.1 |
| *Zea mays* | SPCH1 | Zm00001d046096 | RefGenv4 |
| *Zea mays* | SPCH2a | Zm00001d016095 | RefGenv4 |
| *Zea mays* | SPCH2b | Zm00001d053391 | RefGenv4 |
| *Arabidopsis thaliana* | SPCH | AT5G53210 | Araport11 |
